# Radial contractility of Actomyosin-II rings facilitates cargo trafficking and maintains axonal structural stability following cargo-induced transient axonal expansion

**DOI:** 10.1101/492959

**Authors:** Tong Wang, Wei Li, Sally Martin, Andreas Papadopulos, Anmin Jiang, Golnoosh Shamsollahi, Rumelo Amor, Vanessa Lanoue, Pranesh Padmanabhan, Frederic A. Meunier

## Abstract

Most mammalian neurons have a narrow axon, which constrains the passage of large cargoes such as autophagosomes that can be larger than the axon diameter. Radial axonal expansion must therefore occur to ensure efficient axonal trafficking. In this study we consistently find that the trafficking speed of various large axonal cargoes is significantly slower than that of small ones, and reveal that the transit of diverse-sized cargoes causes an acute, albeit transient axonal radial expansion, which is immediately restored by constitutive contractility. Using live super-resolution microscopy, we demonstrate that actomyosin-II controls axonal radial contractility and local expansion, and that NM-II filaments associate with periodic F-actin rings via their head domains. Pharmacological inhibition of NM-II activity, significantly increases axon diameter by detaching the NM-II from F-actin, and impacts the trafficking speed, directionality, and overall efficiency of long-range retrograde trafficking. Consequently, prolonged disruption of NM-II activity leads to disruption of periodic actin rings and formation of focal axonal swellings, a hallmark of axonal degeneration.

**Summary:** Axonal radial contractility and local expansion control the retrograde trafficking of large cargoes. The periodic actomyosin-II network comprises of NM-II filaments and F-actin rings. Loss of actomyosin-II-mediated radial contractility causes defects in axonal trafficking and stability, leading to degeneration.

## Introduction

Neurons are polarized cells that contain many nerve terminal boutons separated from the cell body by a long and thin axon. Within the axon, an active bidirectional cargo transport system mediates the trafficking of proteins, lipids, membrane-bound vesicles and organelles (cargoes) that undergo retrograde or anterograde transport. Tightly regulated axonal transport is pivotal for neuronal development, communication and survival (Barford et al., 2017; Tojima and Kamiguchi, 2015). Despite the heavy trafficking, quantitative electron microscopy studies have found that thin axons (inner diameter < 1 μm) are the most abundant type in the mammalian central nervous system (CNS) (Liewald et al., 2014; Perge et al., 2012). For instance, the long-range connective axons found in the human corpus callosum have an average diameter that ranges between 0.64 μm and 0.74 μm (Liewald et al., 2014). In contrast, the size of axonal cargoes is highly variable, encompassing autophagosomes (0.5-1.5 μm) (Mizushima et al., 2002), mitochondria (0.75-3 μm) (McBride et al., 2006) and endosomes (50 nm-1 μm) (Altick et al., 2009). Thus, the range of cargo sizes is comparable to, or surprisingly even larger than some of the CNS axons themselves. This advocates for the existence of radial contractility in the axons, which would allow the transient expansion of axon calibre and facilitate the passage of large cargoes. Indeed, the expansion of axonal diameter surrounding large cargoes, *i.e.* autophagosomes (Wang et al., 2015) or mitochondria (Yin et al., 2016), has been observed by super-resolution microscopy and 2D-electron microscopy (EM) in both normal and degenerating axons (Giacci et al., 2018; Maia et al., 2015). Considering the spatial constriction exerted by the rigid and stable circumventing actin MPS (membrane-associated periodic cytoskeletal structures) (Abouelezz et al., 2019a; Qu et al., 2017; Zhang et al., 2017; Zhong et al., 2014), the trafficking of large cargoes is likely to be affected. In fact, a simulation study based on axon structure and intra-axonal microfluidic dynamics predicted that cargo trafficking was impeded by the friction from the axonal walls in small-calibre axons (Wortman et al., 2014). In line with this prediction, correlations between axon diameter and axon trafficking have been recently reported in *Drosophila* (Fan et al., 2017; Narayanareddy et al., 2014) and rodent neurons (Leite et al., 2016; Pesaresi et al., 2015). However, direct evidence showing whether and how axonal radial contractility affects cargo trafficking is still lacking.

We hypothesized that the underlying structural basis for axonal radial contractility is the subcortical actomyosin network, which is organized into specialized structures called membrane-associated periodic cytoskeletal structures (MPS), as revealed with super-resolution microscopy along the shafts of mature axons (Xu et al., 2013). F-actin, together with adducin and spectrin, forms a subcortical lattice with a ∼190 nm periodic interval covering the majority of the axon length (Han et al., 2017; Xu et al., 2013). Disrupting axonal F-actin or spectrin leads to disassembly of MPS (He et al., 2016; Huang et al., 2017; Zhong et al., 2014), which initiates axonal degeneration (Unsain et al., 2018; Wang et al., 2019). In addition, the depletion of adducin causes progressive dilation of the axon diameter and axon loss, accompanied with slightly impaired axonal trafficking (Leite et al., 2016). The fact that adducin knock-out axons are still capable of decreasing the diameter of actin rings over time suggests the existence of additional actin regulatory machineries that maintain this constriction. Indeed, the dynamic contractility of the subcortical actomyosin network depends on non-muscle myosin-II (NM-II) (Even-Ram et al., 2007; Salbreux et al., 2012), which generates the subcellular forces required to restore the shape of the cell following acute stretching (Papadopulos et al., 2015). In line with this function, the activated regulatory light chain (p-MRLC) of NM-II (Berger et al., 2018; Evans et al., 2017), as well as Tropomyosin isoform Tpm 3.1 (Abouelezz et al., 2019b), which activates and recruits NM-II to actin fibres (Bryce et al., 2003; Gateva et al., 2017), has recently been shown to coexist in periodic patterns with the actin MPS to maintain the function and structure of the axon initial segment (AIS) (Berger et al., 2018). Indeed, this actomyosin-dependent contractility is implicated in maintaining axon diameter by coupling the radial and axial axonal contractility in *Drosophila* (Fan et al., 2017). Understanding how the dynamic cytoskeletal architecture coordinates the radial axonal contractility and cargo trafficking is therefore warranted.

In this study, we combined live-imaging confocal microscopy and microfluidic techniques to examine the correlation between the speed of axonal cargoes undergoing long-range transport and their size. We found that the speed inversely correlates with the cargo size. Next, using time-lapse super-resolution structured illumination microscopy (SR-SIM), we found that both the axonal plasma membrane and the underlying actin rings undergo dynamic local deformation during the passage of large cargoes, which promotes a transient expansion immediately followed by constriction. We further demonstrated that this transient change in axon diameter is controlled by NM-II activity, and that NM-II filaments closely associate with periodic actin rings via their heavy chain head domain. Our results suggest that CNS axons are under constitutive radial constriction, which limits their diameter. Accordingly, short-term inhibition of NM-II activity with either blebbistatin (Kovacs et al., 2004) or ML-7 (Saitoh et al., 1987), does not affect the periodicity of actin rings, but effectively decreases their contractility and tilting angle, thereby expanding the axonal diameter. As a result of augmented axon diameter, blebbistatin increases both the speed of directed large cargoes and the back-and-forth movements of undirected ones. This leads to an increase in cargo mobility at the expense of overall trafficking efficacy. Prolonged NM-II inactivation by either NM-II shRNA or transfection of a myosin-II regulatory light chain (MRLC) loss-of-function mutant disrupts the MPS structure and leads to the formation of focal axon swellings (FAS) - a hallmark of axonal degeneration. In conclusion, our study reveals a novel critical role of axonal NM-II which, by associating with subcortical MPS in the axon shaft, provides subcellular radial constriction that minimizes the axonal swelling and undirected cargo movements. It therefore ensures structural stability as well as cargo-trafficking efficiency along the small-calibre axons.

## Results

### The speed of retrograde axonal cargoes is inversely correlated with their size

Large axonal cargoes such as endosomes, lysosomes, autophagosomes and mitochondria tend to accumulate in FAS under pathological conditions (Tammineni et al., 2017), suggesting that the size of cargoes might alter the axonal trafficking efficacy. To determine the relationship between the size of cargoes and their transport speed, we analyzed the speeds of various-sized retrograde lysosomal and endosomal vesicles. These cargoes were generated and fluorescently labelled with the lysosomal marker Lysotracker or the endosomal marker Cholera toxin subunit B (CTB) at the nerve terminals, and underwent retrograde trafficking along the axon bundles of live hippocampal neurons cultured in microfluidic devices (Fig. 1a, b). Hydrostatic pressure was used to restrict the labelling reagents to the terminal chamber during the 5 min pulse-labelling (Fig. 1c). This was followed by a thorough wash in culture medium to remove the excess fluorescent probe and confocal time-lapse imaging and automatic tracing of the fluorescently tagged cargoes as previously described (Joensuu et al., 2017; Wang et al., 2016; Wang et al., 2015). To investigate different stages of axonal trafficking, live-imaging was performed in two distinct axonal regions: (1) the axon shafts adjacent to the soma chamber (Fig. 1d, left) and (2) within the terminal chamber (Fig. 1d, right). Lysotracker-labelled vesicles detected in the nerve terminal chamber exhibited very confined movements (sFig. 1a and sVideo 1). We further quantified the average speed of these tracks, which were sorted into two different groups according to their diameter (sFig. 1a, bottom panels). We observed that vesicles with a large diameter (‘large’, diameter > 0.5 μm) moved significantly slower (0.115 ± 0.039 μm/s) than those with a smaller diameter (‘small’, diameter 0.5 μm; 0.159 ± 0.006 μm/s), as shown in sFig. 1b. This suggests that transport of large axonal cargoes in axons surrounding nerve terminals could be impeded during their transit. To specifically investigate the correlation between the size of axonal cargoes and their active transport speed, we further examined the trafficking speed of long-range CTB-positive retrograde carriers in the soma-proximal axon channels (Fig. 1e-g). Consistent with our previous study (Joensuu et al., 2016), these long-range carriers exhibited a much greater trafficking speed (0.974-1.659 μm/s) than the carriers at nerve terminals (0.115-0.159 μm/s; Fig. 1g; sFig. 1b). Similar to the lysosomes in the nerve terminals, the trafficking speed of these CTB-positive carriers also inversely correlated with their diameter, with small-diameter cargoes moving faster (1.657 ± 0.06 μm/s) than large-diameter ones (0.974 ± 0.05 μm/s; Fig. 1g; sVideo 2). We then examined the correlation between cargo size and speed by plotting the apparent diameter of either lysosomal carriers or CTB-positive carriers against their speed. This revealed a negative correlation, with a Pearson’s coefficient of -0.303 ± 0.095 and -0.273 ± 0.036 between cargo diameter and trafficking speed in the terminal (“Lysotracker”, Fig. 1h) and proximal (“CTB”, Fig. 1h) axons, respectively, indicating that the speed of the trafficked cargoes declines as the cargo size increases.

**Figure 1.**
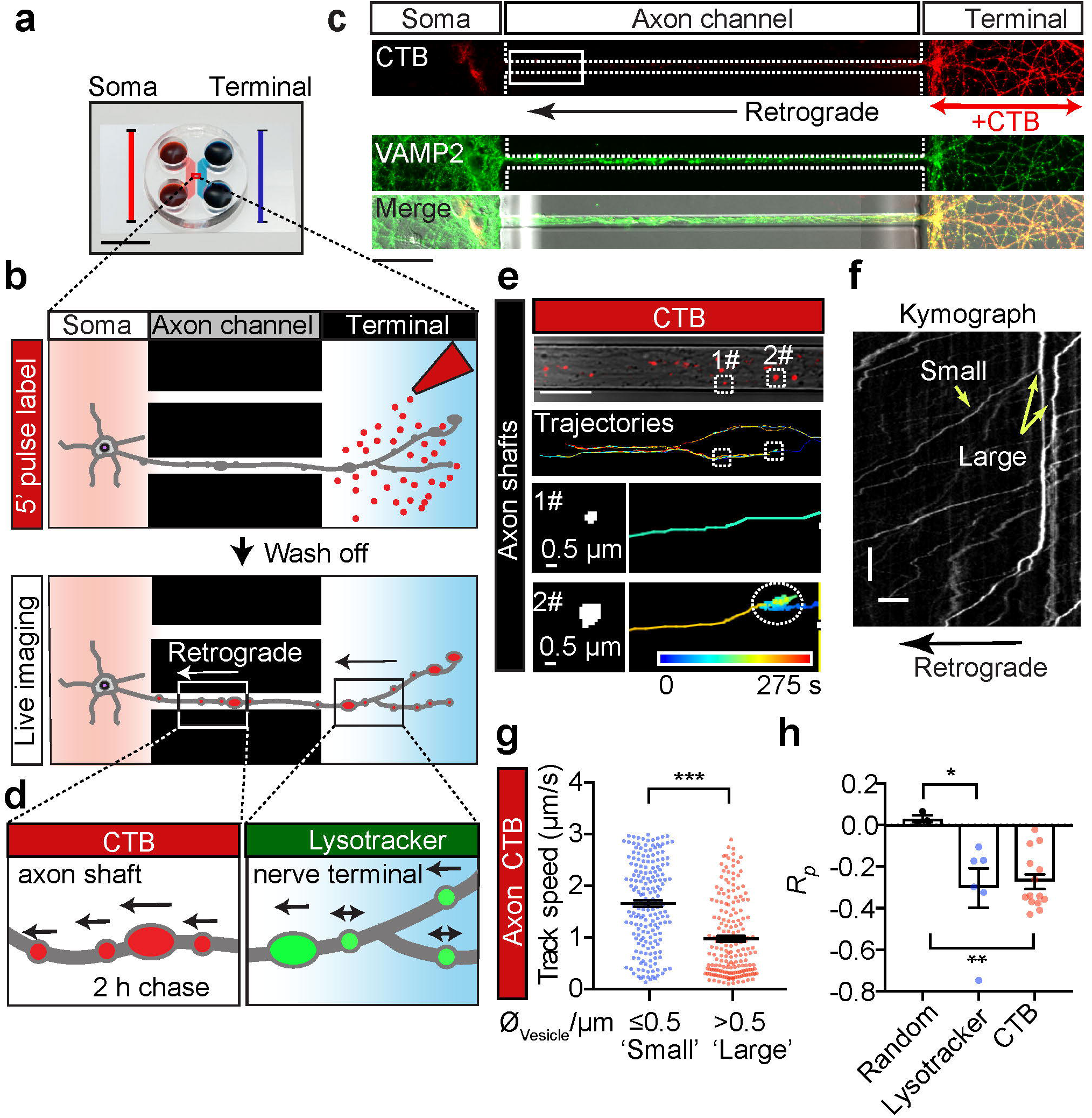
The speed of retrograde axonal transport cargoes is inversely correlated with their size. **(a)** Microfluidic chambers isolate unidirectional axon bundles, bar =1 cm (adapted from Xonamicrofluidics.com). (**b)** Schematic diagram of the pulse-chase labelling process. Cultured hippocampal neurons were grown in a microfluidic device for 14 days *in vitro* (DIV). The nerve terminal chamber was incubated with fluorescently tagged CTB (50 ng/ml) for 5 min or Lysotracker Deep Red (50 nM) for 30 min (pulse). After thorough washes and a 2h chase. (**c)** Representative images of cultured neurons, showing the restriction of retrograde CTB surface labelling to nerve terminals and the position of the observation window (white box). Scale bar = 50 µm. **(d)** The axonal retrograde transport of CTB or Lysotracker was monitored, at the level of the proximal axon shafts or in the nerve terminal chamber, respectively. (**e)** Time-lapse images of CTB carriers. Top panels: CTB labelling and tracing trajectories within the axon channels. Trajectories of small (#1, diameter 0.5 μm) and large (#2, diameter > 0.5 μm) carriers are magnified in the bottom panels, respectively. (**f)** Representative kymographs of CTB-positive cargoes along a single axon, depicting track displacements of small and large carriers. x-bar = 10 μm; y-bar = 10 s. (**g)** Grouped analysis of the average speeds of CTB cargoes with small (diameter 0.5 μm) and large (diameter > 0.5 μm) diameters. Data represent mean ± s.e.m (small, n=187, large n=185 tracks from 3 independent preparations; two-tailed unpaired t-test, \*\*\**p*<0.001). (**h)** Pearson’s coefficient of the speed and diameter of retrograde Lysotracker-positive and CTB-positive cargoes. Data represent mean ± s.e.m from 3 independent preparations (random, n=3 simulated data sets; Lysotracker, n=6; CTB, n=14; n represents the number of axon channels analysed; the single value of the average correlation coefficient between the size and speed of all trajectories in each axon channel was calculated and used for the plot. 3 Independent groups of Gaussian-distributed random numbers were generated using the *normrnd* function of Matlab. Two-tailed unpaired t-test, **p<0.01, *p<0.05).

### Transit of large cargoes causes a significant transient radial expansion of the axonal plasma membrane and the underneath periodic actin rings

Each organelle undergoing retrograde axonal transport is driven by multiple dyneins, which are stochastically activated and collectively drive cargo transport through the axonal cytosol (Chowdary et al., 2015b; Mallik et al., 2005; Rai et al., 2013). Given the low viscosity of axonal cytosol, the force generated by cooperative dyneins is sufficient to ensure their retrograde trafficking through the axon (Chowdary et al., 2015b). Thus, the reduced speed of the larger cargoes we observed is unlikely due to insufficient driving force or a higher viscous load due to the larger size. Considering the recent evidence suggesting the role of axon diameter in axon cargo trafficking (Fan et al., 2017; Leite et al., 2016; Narayanareddy et al., 2014), we hypothesized that the size-dependent friction on the axonal cargoes comes from the constrictive force exerted by the axonal plasma membrane, which is more likely to impede the transport of larger retrograde cargoes.

To test this hypothesis, we examined the diameter of axons in the presence or absence of cargoes at the ultrastructural level. In order to eliminate confounding factors related to the analysis of dendrites, experiments were only performed on axonal bundles formed within the channels of microfluidic devices (Joensuu et al., 2017; Wang et al., 2016; Wang et al., 2015), as shown in Fig. 2a. We first used EM to visualize the morphology of both axon shafts and their internal cargoes. As demonstrated in Fig. 2b-c, on the parallel axonal bundles, the diameters of axons were indeed significantly increased around large cargoes, such as large endosomes (Fig. 2c, i, arrow), mitochondria (Fig. 2c, iii, white arrowheads) and autophagosomes (Fig. 2c, iii-iv, black arrowheads). When we measured the diameter of the axonal segment with (red) and without cargo (blue) in the same axon, we found that those with cargoes had a significantly larger diameter (347 ± 15.6 nm) than those without cargoes (259 ± 9.4 nm; Fig. 2d) with paired comparison. We also observed that, as the size of the cargoes increased, the extent of axon expansion also increased proportionally (Fig. 2e, f), suggesting that the stretch of the axon membrane is indeed caused by the transiting cargo.

**Figure 2.**
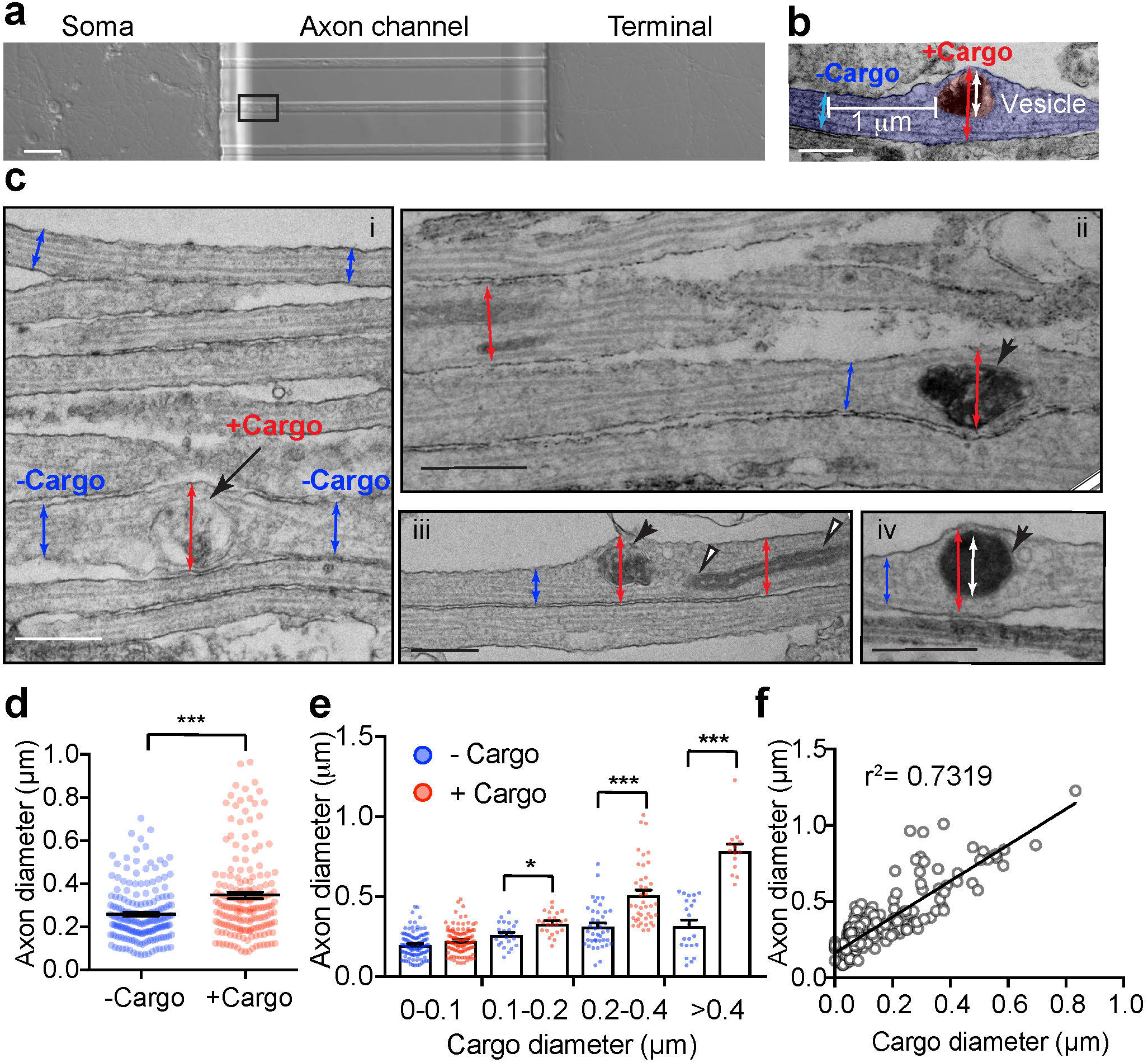
The size of axonal cargoes correlates with the diameter of the axon. **(a)** Brightfield image of DIV14 rat hippocampal neurons cultured in a microfluidic device with the region selected for EM outlined. Scale bar = 250 μm. (**b)** Representative electron micrographs showing the axonal diameter measurements with cargo (red) or without cargo (blue), and the associated cargo size (white). Scale bar = 0.5 μm **(c)** Electron micrographs of axon bundles from hippocampal neurons cultured in microfluidic devices. (**i-iv)** Axon diameters with and without cargo are marked with red and blue arrows, respectively. (**i)** Endosome = black arrow, (**i-iii)** Mitochondria = white arrowheads, autophagosome = black arrowheads. (**iv)** Inner diameters of cargoes are marked with white arrows. Bars = 500 nm. (**d)** Quantification of axon diameters with cargo and without cargo. Data represent mean ± s.e.m, n= 182 (+ Cargo) and 182 (- Cargo) measurements from 2 independent preparations (two-tailed paired t-test, \*\*\**p*<0.001). (**e)** Grouped quantification of axonal diameter as a function of binned vesicle size. Data represent mean ± s.e.m; for the + Cargo group from left to right: n=104, 23, 41 and 14 measurements; for the - Cargo group from left to right: n=104, 23, 41 and 22 measurements, data were from 2 independent preparations (two-tailed unpaired t-test, \**p*<0.05; \*\*\**p*<0.001). (**f)** Cross-correlation analysis of cargo size and axonal diameter. Linear regressions were performed with the 182 paired measurements of axonal and cargo diameters of the (+ Cargo) group, data from 2 independent preparations.

We next investigated the effect of transiting cargoes on the diameter of axons in live hippocampal neurons. To effectively label the subcortical actomyosin network in axons, we used Lifeact-GFP, a peptide that binds to both actin filaments (F-actin) and cytosolic actin monomers (Riedl et al., 2008). Similar to a previous study (Ganguly et al., 2015), with the resolution of SIM, we detected Lifeact-GFP distribution in both the filamentous and cytosolic fractions in axons of live hippocampal neurons, we also observe various-sized intra-axonal fluorescence voids that likely represent axoplasmic organelles (Fig. 3a), which are known to exclude actin from their lumens (Gormal et al., 2015). Similar to the cargo-induced axon dilation observed with EM, the axon diameter was also significantly expanded in axon segments containing these unlabelled organelles (Fig. 3b). We then characterized the nature of these organelles by comparing their localization with that of various organelle markers resolved by 3D-SIM, and found substantial overlap with autophagosomes (LC3-mRFP, sFig. 2a), late endosomes (Rab7-mRFP, sFig. 2b) and mitochondria (Mito-TagRFP, sFig. 2c). This suggests that these organelles caused significant local dilation of the axon (sFig. 2d). To further investigate whether these organelles were cargoes that associate with the retrograde transport machinery, we determined their colocalization with markers of retrograde carriers, such as terminal-derived CTB (Wang et al., 2016) and the neuron-specific dynein intermediate chain 1B (DIC^1B^) (Ha et al., 2008). These organelles partially overlapped with CTB- and DIC^1B^-positive axonal structures (sFig. 2e, f), indicating that a substantial portion of them were indeed caused by retrograde trafficking organelles in live axons.

**Figure 3.**
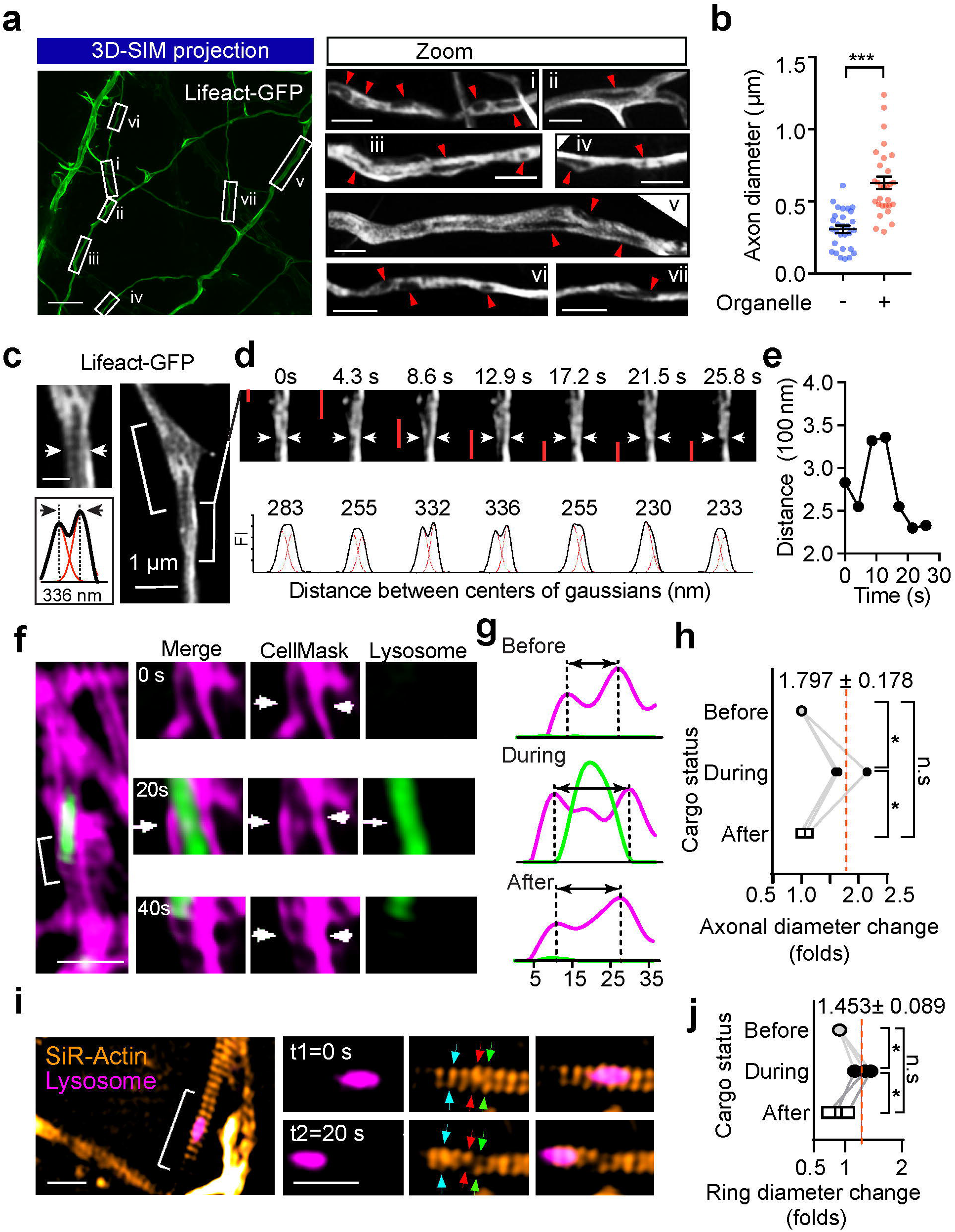
The passage of large axonal cargoes causes a transient radial expansion of the axon. **(a)** Rat hippocampal neurons were transfected with Lifeact-GFP and imaged with 3D-SIM. Left: a representative maximum projection of 3D-SIM of Lifeact-GFP expressing axons are shown; Bar = 5 μm. Right: magnified regions of interest (ROIs) in left panel, arrowheads indicate fluorescence voids with low Lifeact-GFP signals within the axon. Bar = 1μm. **(b)** Quantification of axon diameters with (+) and without (-) unlabelled organelles. Data represent mean ± s.e.m from 3 independent preparations (+ black-hole, n=29, - black-hole, n=29 axons; two-tailed unpaired *t*-test, \*\*\**p*<0.001). **(c)** Rat hippocampal neurons cultured in a glass-bottom dish were transfected with Lifeact- GFP (DIV 12) and imaged by time-lapse SIM (DIV 14). Representative live axons with unlabelled cargos passing through are shown, with inset demonstrating the Gaussian fittings of the annotated line transection of axon. Bar = 0.5 μm (inset). **(d)** Time-lapse images of bracketed region in **(c)**, showing the axonal diameter fluctuation as the cargo (indicated with red bar) transits. **(e)** Plot of the distance between axon membranes against time. **(f)** Representative time-lapse dual-colour SIM of live axons with plasma membrane labelled with CellMask and Lysosome with Lysotracker-Red. The deformations of plasma membrane trigged by the passage of lysosome (arrows) were indicated with arrowheads. Bar = 1 μm. **(g)** Line transection of axon plasma membrane annotated with blue arrows. **(h)** Quantification of axonal diameter changes as cargoes pass through. Data represent mean ± s.e.m from 3 axons (two-tailed paired *t*-test, \**p*<0.05; n.s. no significant difference). **(i)** Representative time-lapse dual-colour SIM of live axons with periodic actin rings labelled with SiR-Actin and Lysosome with Lysotracker-Red. The deformations of actin rings trigged by the passage of lysosome were indicated with arrowheads. Bar = 1 μm. **(j)** Quantification of actin ring diameter changes as cargoes pass through. Data represent mean ± s.e.m from 3 axons (two-tailed paired *t*-test, \**p*<0.05; n.s. no significant difference).

Next, we investigated whether the transit of cargoes correlated with the local axon dilation in live neurons, using time-lapse SIM. In Lifeact-GFP expressing axons, using unbiased Gaussian fitting (Fig. 3c), we assessed the fluctuations in axon diameter and clearly detected radial diameter expansion through the transient separation of the two lateral axonal membranes, which caused an increase in the distance between the centre of the Gaussians (Fig. 3d; sVideo 3). This effect was transient and the initial diameter was restored after the passage of the unlabelled organelles, as shown in Fig. 3e. Similarly, we used CellMask to label the axonal plasma membrane, and found that an axonal diameter expansion occurred concomitantly with the passage of Lysotracker-positive cargoes (Fig. 3f-h; sVideo 4, 5). Interestingly, this cargo-induced radial expansion was also observed in the SiR-Actin labelled periodic actin rings (Fig. 3i-j; sVideo 6, 7). Taken together, our results demonstrate that passage of axonal cargoes could cause transient radial expansion of the axonal diameter, including both plasma membrane and underlying periodic actin rings in live neurons.

### NM-II controls the radial contractility of the periodic actin rings along the axon

The contractility and plasma membrane tension is controlled by the actomyosin-II network, which is composed of NM-II filaments sliding upon filamentous actin (F-Actin) (Arnold and Gallo, 2014; Berger et al., 2018; Evans et al., 2017). This interaction can be disrupted by blebbistatin, a specific membrane-permeable inhibitor that blocks the ATPase activity of the myosin heavy chain (MHC) and detaches NM-II from F-actin (Kovacs et al., 2004), as shown in Fig. 4a. To explore the molecular basis of the axonal radial contractility, we first sought to resolve the actomyosin structures along the axon shafts. To specifically label actin MPS in live neurons, we employed SiR-Actin, a far-red fluorescent probe that has high brightness and low cytotoxicity (Lukinavicius et al., 2014; Lukinavicius et al., 2013) to label the F-actin along the axon. We used time-lapse SIM, which had previously been used to accurately visualize axon MPS in live neurons (Qu et al., 2017), to visualize the periodic axon actin rings along live axons (Fig. 4b; sVideo 8). We found MPS formed along the axon shaft has a conserved longitudinal spacing of ∼190 nm (193.2 ± 0.15 nm, Fig. 4c), which is similar to the spacing distance observed in fixed and Phalloidin-647-labelled axons (191.8 ± 2.3 nm; sFig. 3a-d). These values were similar to those previously reported in rat hippocampal axons (He et al., 2016; Xu et al., 2013). As NM-II is involved in the regulation of axonal diameter (Fan et al., 2017) and associated with the MPS in axons (Berger et al., 2018), we also examined whether blocking NM-II activity using blebbistatin would affect the spacing of actin MPS. We found that 60 min of blebbistatin treatment (10 μM) had no effect on the spacing (Fig. 4c; sFig. 3a-e). However, this short-term NM-II inactivation significantly increased the radial diameters of the actin rings in treated axons (Fig. 4d; sFig. 3f), and also decreased the contractility of actin MPS, as reflected in the decreased ring diameter fluctuations over 20 s in live axons (Fig. 4e). Similar effects on axonal diameter expansion were also observed in fixed axons labelled using Phalloidin-647 (sFig. 4a, b). Moreover, the effect of blebbistatin was reversible, as reflected by the partially recovered diameter of periodic actin rings following blebbistatin washout in SiR-Actin-labelled live axons (sFig. 4c, d). In addition, the viability of treated neurons was unaffected by blebbistatin treatment (sFig. 4e, f). The significant dilation of periodic actin rings caused by detaching NM-II suggests that the interaction between NM-II and periodic actin rings controls the radial contractility of axons. Interestingly, we found that blebbistatin treatment also affected the orientation of actin rings by decreasing their tilting angles with the axonal axis (Fig. 4f). This suggests NM-II is likely to exert tension in-between adjacent actin rings.

**Figure 4.**
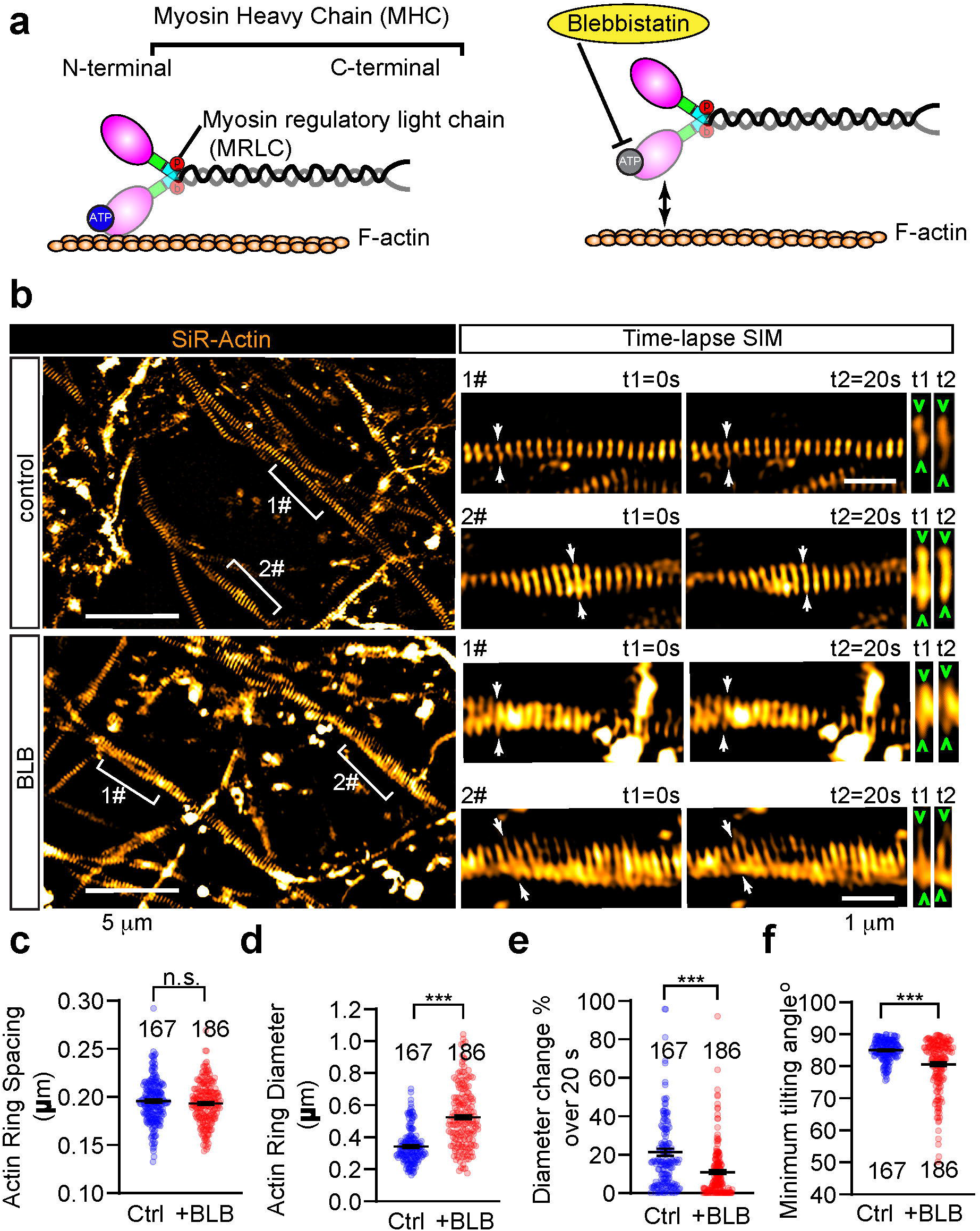
Short-term inactivation of NM-II affects the diameter and angle of axonal actin rings, but not their periodic spacing. **(a)** Cartoon showing the organization of actomyosin structure in non-muscle cells, with the ATP binding-site in the head domain of MHC annotated. On the treatment with blebbistatin, ATPase activity of NM-II is blocked, leading to its detachment from the F-Actin. **(b)** In cultured hippocampal neurons, endogenous periodic axonal actin rings were labelled using SiR-Actin and live-imaged using 2D-SIM. Representative time-lapse SIM images of axonal actin rings are shown before (control) and after short-term blebbistatin treatment (10 µM, 30-60 min). Bracketed regions are magnified in right panels. Dynamic diameter changes of actin rings are annotated with arrowheads. Bar = 5 µm (left) and 1 µm (right). **(c-f)** Quantification of the spacing **(c)**, diameter **(d)**, fluctuation of actin ring diameter **(e)** and minimum tilting angles **(f)** of the periodic actin rings along the axon. Data represent mean ± s.e.m. N values are labelled on the panels, representing numbers of axonal actin rings analysed. Values were measured from 3 independent cultures (two-tailed unpaired *t*-test, \*\*\**p*<0.001; n.s. no significant difference.

As shown in Fig. 5a, the local contractile activity of NM-II was also controlled by the myosin regulation light chain (MRLC), which is phosphorylated on conserved sites and is critical for the formation of periodic actin rings in the AIS (Berger et al., 2018). To further explore the activity of NM-II on axon radial contractility, we used ML-7 and Calyculin A, which inhibits and activates the NM-II respectively by controlling MRLC phosphorylation levels (Kato et al., 1988; Saitoh et al., 1987). The efficacy of ML-7 and Calyculin A treatment was confirmed by western blotting using specific MRLC diphosphorylation antibody (p-MRLC). Blebbistatin, the MHC ATPase inhibitor (Kovacs et al., 2004), has no effect on the p-MRLC level (Fig. 5b). Along the axons of SiR-Actin labeled live neurons, we found that ML-7 treatment (10 μM, 30 min), which inhibits NM-II activity, significantly expanded the diameter of periodic actin rings (Fig. 5c, d) – an effect similar to that of blebbistatin. To our surprise, Calyculin A treatment (50 nM, 30 min), which activates NM-II, had no significant effect on the diameter of periodic actin rings (Fig. 5d). Both ML-7 and blebbistatin affected the actin ring tilting angle (Fig. 5e), but not their spacing (Fig. 5f). To further investigate the correlation between p-MRLC and axonal diameter, we also examined the correlation between the endogenous p-MRLC level and axon diameter (sFig. 5a). We found no significant correlation (sFig. 5b). These data show that raising NM-II activity by targeting MRLC does not change the diameter of actin rings, whereas inhibiting NM-II activity significantly increases their diameter, indicating that axonal NM-II is constitutively activated, which is likely to exert a constant tension on the abutting axon plasma membrane.

**Figure 5.**
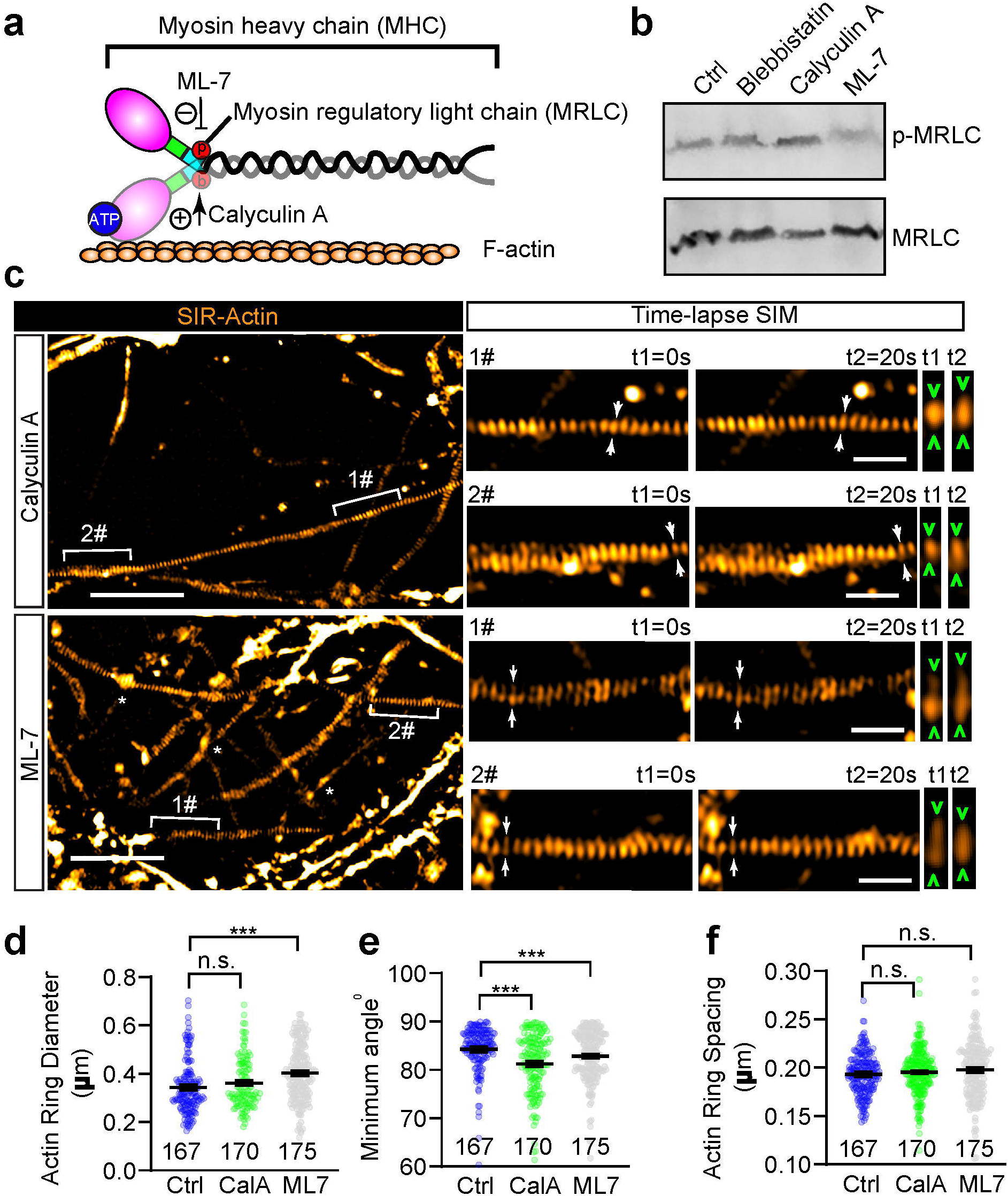
Inhibition of MRLC phosphorylation slightly affects the diameter and tilting angle of axonal actin rings, but not their periodic spacing. **(a)** Cartoon showing the organization of actomyosin structure in non-muscle cells, with the effecting sites of ML-7 and Calyculin A on MRLC annotated. **(b)** Western blot showing the level of diphosphorylated MRLC (p-MRLC) following 30 min treatment of Blebbistatin (10 μM), ML-7 (10 μM) and Calyculin A (50 nM). **(c)** In cultured hippocampal neurons, endogenous periodic axonal actin rings were labelled using SiR-Actin and live-imaged using 2D-SIM. Representative time-lapse SIM images of axonal actin rings are shown following 30 min of treatment with ML-7 (10 μM) or Calyculin A (50 nM) respectively. Bracketed regions are magnified in right panels. Dynamic diameter changes of actin rings are annotated with arrowheads. Bar = 5 µm (left) and 1 µm (right). **(d-f)** Quantification of the diameter **(d)**, minimum tilting angle **(e)** and spacing **(f)** of the periodic actin rings along the axon. Data represent mean ± s.e.m. N values are labelled on the panels, representing numbers of axonal actin rings analysed. Values were measured from 3 independent cultures (two-tailed unpaired *t*-test, \*\*\**p*<0.001; n.s. no significant difference).

### NM-II closely associates with periodic actin rings along the axon

To assess the relationship between NM-II and periodic actin rings along the axon shaft, we used dual-colour 3D-SIM to investigate the distribution of endogenous NM-II and actin rings. We co-labeled the axons of DIV14-28 rat hippocampal neuron using SiR-Actin or Phalloidin and an antibody that recognizes either the N-terminus head domain (αNM-II(nt)) or the C-terminus rod domain of NM-IIB (αNM-II(ct)), which is the dominant form of NM-II in mature axons (Berger et al., 2018) (Fig. 6a). Following fixation and permeabilization, the number of axonal segments displaying periodic actin rings was reduced (Fig. 6b, c). We therefore restricted our analysis of NM-II and actin dual labeling to the axonal segments with preserved periodic actin rings.

**Figure 6.**
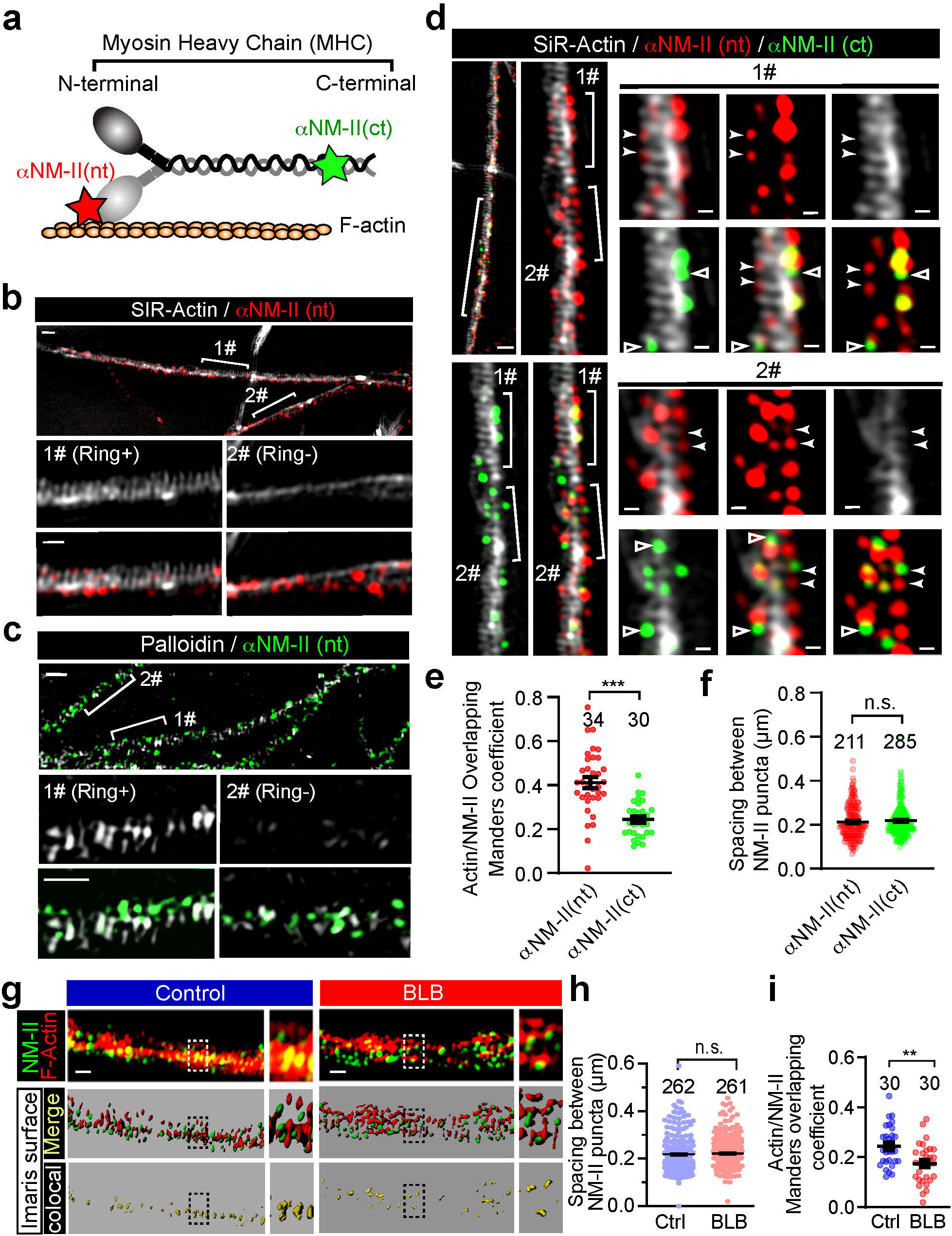
Periodic actin rings correlate more extensively with the head domain than the rod domain of the NM-II filaments along axons. **(a)** Cartoon showing the binding site of antibodies against the NM-II head domain (αNM-II (nt)) and rod domain (αNM-II (ct)), respectively. **(b)** After fixation DIV14 rat hippocampal neurons were stained for endogenous F-actin (Phalloidin) and head domain (αNM-II (nt)), and imaged with dual-colour SIM, bracketed regions with persistent actin rings (Ring+) or without actin rings (Ring-) are magnified in the bottom panels, respectively. Bar = 1 μm (top) and 500 nm (bottom). **(c)** Prior to fixation DIV14 rat hippocampal neurons were labelled with SiR-Actin, then were fixed and stained for endogenous head domain (αNM-II (nt)), and imaged with dual-colour SIM, bracketed regions with persistent actin rings (Ring+) or without actin rings (Ring-) are magnified in the bottom panels, respectively. Bar = 1 μm (top) and 500 nm (bottom). **(d)** Triple-colour SIM image of endogenous F-actin (SiR-Actin), head domain (αNM-II (nt)) and rod domain (αNM-II (ct) of the NM-II along the axon of hippocampal neuron. Bracketed regions are magnified on the right. αNM-II (nt) puncta overlapping with actin rings are annotated with arrowheads. αNM-II (ct) puncta not overlapping with actin rings are annotated with hollow arrows. Bar = 1 μm (left) and 200 nm (right). **(e)** Comparison of co-localization between actin rings with either the head domain (αNM-II (nt)) or rod domain (αNM-II (ct)) of NM-II filaments. **(f)** Quantification of the spacing between adjacent NM-II puncta in the axonal segments with preserved periodic actin rings. **(g)** The NM-II and actin structures resolved with 3D-SIM (top) were rendered into surface (middle) using Imaris software; boxed regions are magnified to show the accuracy of the rendering. The colocalization of the bracketed region is shown in the bottom panels. Comparison of colocalization between NM-II and actin rings before and after the 60 min blebbistatin treatment. Bar= 0.5 µm. **(h)** Quantification of the spacing between adjacent NM-II puncta in the axonal segments with preserved periodic actin rings, in control and blebbistatin treated neurons. Data represent mean ± s.e.m. N values are labelled on the panels, representing numbers of axonal actin rings analysed. **(i)** The Mander’s coefficient reflecting the colocalization rate. Data represent mean ± s.e.m, n=30 (control), 30 (BLB) axon segments for fraction of the F-actin overlapping with NM-II; data were from 3 independent cultures (two-tailed unpaired t-test, \*\**p*<0.01). Values were measured from 3 independent cultures (two-tailed unpaired *t*-test, \*\*\**p*<0.001; n.s. no significant difference).

Using the αNM-II(nt), which detects the head domains of NM-II filaments (Fig. 6a), we observed an extensive colocalization between the αNM-II(nt) puncta and periodic actin rings labelled with SiR-Actin (Fig. 6d). In contrast, the αNM-II(ct) (rod domain of NM-II filaments) immunostaining (Fig. 6a), displayed less overlap with the periodic actin rings (Fig. 6d), as compared to αNM-II(nt) (Fig. 6e). We also quantified the spacing between adjacent NM-II puncta along the longitudinal axis of the axon, and found that both αNM-II(ct)- and αNM-II(nt)-stained puncta exhibited a periodic distribution (Arrowheads in Fig. 6d), with an average spacing of ∼200 nm (212.9 ± 5.27 nm, αNM-II(nt); 218.3 ± 4.13 nm, αNM-II(nt); in Fig. 6f). These values are similar to the periodicity of the actin rings (Xu et al., 2013), and also consistent with that of phosphorylated-MRLC periodicity, as reported recently (Berger et al., 2018).

We next investigated whether the distribution pattern of NM-II was affected by the inhibition of its activity, and found that short-term blebbistatin treatment did not significantly alter the NM-II periodicity (Fig. 6g, h), but significantly decreased the degree of colocalisation between NM-II and actin MPS (Fig. 6i). This was most apparent with the reduced colocalization between NM-II and actin voxels, as rendered by Imaris surface function of 3D-SIM images following blebbistatin treatment (sFig6. a, b). Indeed, NM-II and actin rings distributed more discretely from each other in blebbistatin-treated axons (Fig. 6g, left panels), as expected due to NM-II detachment from actin rings following blebbistatin inhibition.

To confirm that the NM-II periodic distribution indeed correlates with actin MPS, we performed Triton X-100 extraction prior to fixation in order to specifically remove the subcortical actin MPS components as previously reported (Zhong et al., 2014). Following this extraction, we observed a significant reduction in actin MPS (sFig. 6c, d), confirming the disruption of the membrane-associated actin MPS. Together with this reduction, we detected a dramatic reduction in NM-II positive puncta (sFig. 6e). This concomitant decrease in both actin MPS and NM-II (sFig. 6f) further supports the notion that NM-II associates with actin MPS. Together, these results suggest that NM-II activity keeps the subcortical periodic actin rings constitutively contracted.

### Short-term inhibition of NM-II activity causes axon dilation and interferes with the long-range retrograde trafficking of large cargoes

To test whether axon radial contractility has a functional role in cargo transport, we examined the effect of blebbistatin on retrograde axonal trafficking in neurons grown in a 6-well microfluidic device (sFig. 7a), where we could restrict the action of blebbistatin to the axon segments by adding it through the middle chamber (sFig. 7b). Widespread dilation of the axonal diameter (sFig. 7c, d) and volume (sFig. 7e) along the longitudinal axis was observed using 3D-SIM following 90 min of blebbistatin incubation. To test the integrity of these treated axons, we further examined the axonal microtubule structure by dual-color SIM using a β-tubulin III antibody and phalloidin (sFig. 7f). We found that neither the microtubule bundle intensity (sFig. 7g) nor width (sFig. 7h) were affected by 60 min blebbistatin treatment. This blebbistatin treatment also failed to affect the mitochondrial anchoring, as the movement of the immobile mitochondrial fraction was not affected (sFig. 7i, j), whereas the mobile fractions with the higher average speeds were significantly increased by the blebbistatin movement (sFig. 7i, j), suggesting that the NM-II-dependent axonal contractility is likely to affect the cargo transport.

**Figure 7.**
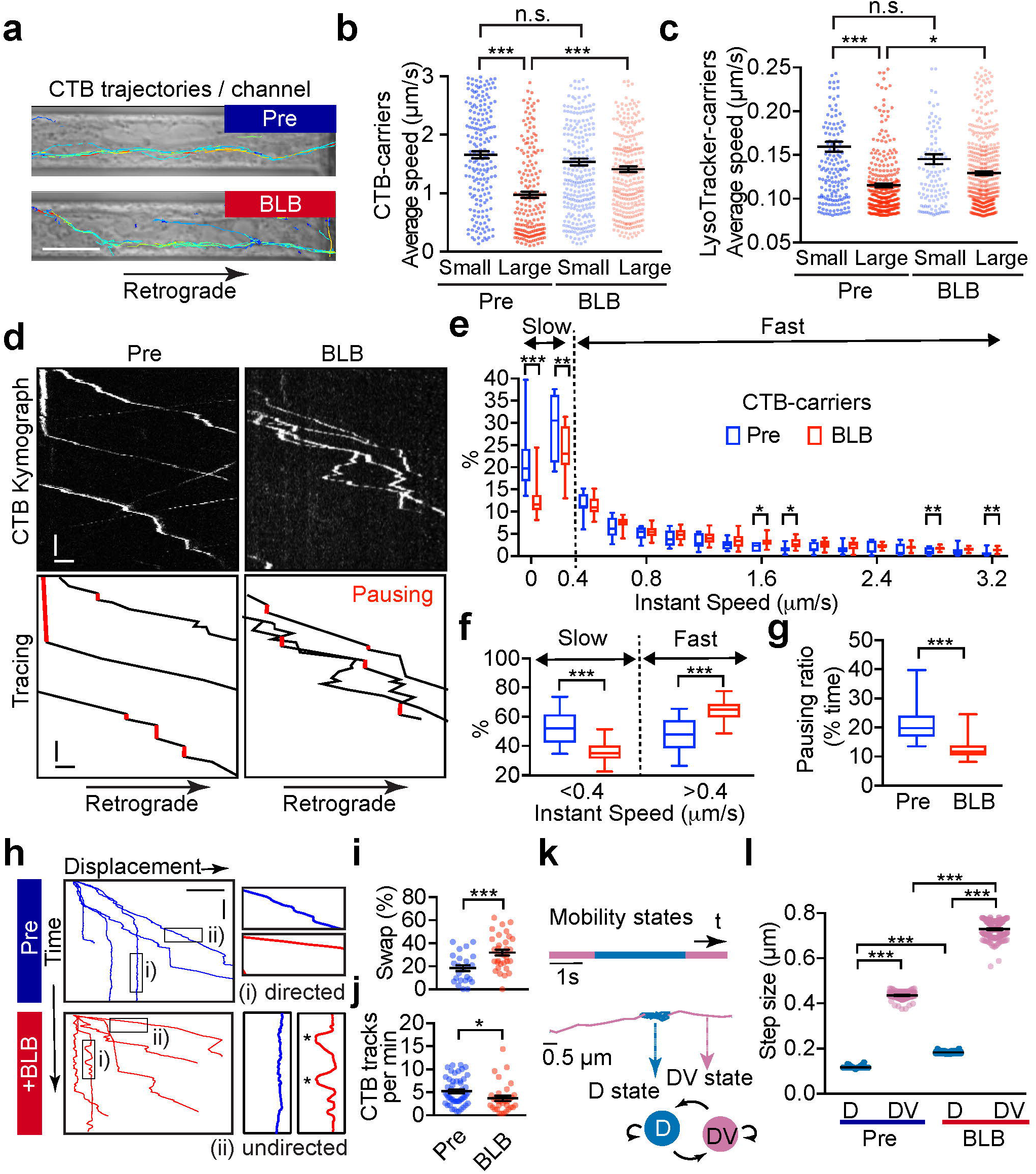
Short-term inactivation of actomyosin-II reduces the efficiency of retrograde axonal trafficking. **(a)** DIV 14 rat hippocampal neurons were cultured in microfluidic devices, and the axon segments were subjected to short-term blebbistatin treatment (10 µM, 60 min). Trajectories of CTB-positive cargoes in the axon channels were traced before (pre) and after (+BLB) blebbistatin treatment. Bar = 10 µm. (**b, c)** Track speed (trajectory length/duration) of the CTB carriers (**b)** or Lysotracker carriers (**c)** before (pre) and after (+BLB) blebbistatin treatment. The speeds of these carriers were sub-grouped according to their diameters. Data represent mean ± s.e.m, n=187, 185, 271, 286 (CTB) and n=179, 342, 128, 573 (Lysotracker) trajectories from 3 independent cultures (two-tailed unpaired t-test, \**p*<0.05, \*\*\**p*<0.001, n.s. no significant difference). The same data of pre-treated groups were used in Figure 1G and Figure S1B. **(d)** Representative kymographs of CTB-carriers in single axons before (Pre) and after 60-min blebbistatin (+BLB) treatment. Tracings of retrograde CTB carriers are shown in the right panels. Pausing states with the trajectories are indicated with red. xbar = 5 s, ybar=1 µm. **(e)** Frequency distribution of the instant speed of CTB-carriers. Showing a significant decrease in the frequency of slow carriers (0-0.4 μm/s) and a significant increase in the fast carriers (0.4-3.2 μm/s). **(f)** Grouped analysis of instant speeds of slow or fast CTB carries, showing a significant difference before (Pre) and after 60-min blebbistatin (+BLB) treatment. **(g)** The ratio of pausing CTB-carriers (instant speed 0-0.2 μm/s). Data are shown in box and whiskers (Min to Max), n=11 (Pre) and 19 (+BLB) axon channels from 3 independent preparations (two-tailed unpaired t-test, \**p*<0.05; \**p*<0.01; \*\*\**p*<0.001) for **e-g**. (**h)** Displacement-time plot of representative Imaris-traced CTB trajectories. (i) Directed, and (ii) Undirected state, as magnified on the right, with the back-and-forth movements marked with asterisks. x-bar = 10 µm, y-bar= 20 s. (**i)** Time ratio of cargo travelling in the reverse direction (swap) to total time travelled. Data represent mean ± s.e.m, n=28 (Pre) and 34 (+BLB) channels from 3 independent preparations (two-tailed unpaired t-test, \*\*\**p*<0.001). (**j)** Quantification of the frequency of CTB-labelled vesicles that cross the observation window per minute. Data represent mean ± s.e.m, n = 51 (Pre) and 32 (+BLB 60’) channels from 3 independent preparations (two-tailed unpaired t-test, \**p*<0.05). (**k)** CTB trajectory displays directed (D, blue) and undirected (DV, pink) motion states inferred by HMM-Bayes analysis. Example of an annotated trajectory colour-coded with the indicated motion states. The time line shows the temporal (s) sequence of the inferred D and DV motion states. (**l)** Step sizes of the two motion states before (pre) and after (+BLB) blebbistatin treatment. Data represent mean ± s.e.m, from left to right, n= 126, 126, 190, 190 different trajectories from 3 independent preparations (two-tailed unpaired t-test, \*\*\**p*<0.001).

To further assess the effects of axonal contractility on cargo trafficking, we examined the impact of disrupting NM-II activity on retrograde trafficking in short-term blebbistatin-treated axons. The average speed of CTB-positive carriers before (pre) and after (BLB) blebbistatin treatment for 60 min (Fig. 7a) were compared. As demonstrated earlier (Fig. 1g), small carriers moved faster than large ones in the absence of blebbistatin (Fig. 7b), whereas blebbistatin treatment specifically increased the trafficking speed of large CTB-positive carriers, but not that of small ones (Fig. 7b and sVideo 9). Similarly, an increase of the average speed was observed for large Lysotracker-positive carriers, but not for small ones (Fig. 7c, sVideo 10). These results indicate that the short-term relaxation of the axonal actomyosin-II network has an initial positive impact on the trafficking of large cargoes, suggesting that axon radial contractility exerts a local brake on their transport.

Given that the large CTB-positive carrier population showed the most significant increase in speed upon blebbistatin treatment (Fig. 7b), we chose these carriers for more detailed motion analyses, in order to further dissect the impact of radial contractility on axonal transport. We used kymograph and instantaneous speed to analyze the movement of individual CTB-carriers. We found that 60 min treatment with BLB decreased the proportion of time that individual carriers spent in pausing (Fig. 7d, red lines; quantified in Fig. 7g) and slow-moving state (instantaneous speed 0-0.4 μm/s) (Fig. 7e, ‘slow’). Concomitantly, the ratio of time spent in fast-moving (instantaneous speed 0.4-3.2 μm/s) was significantly increased (Fig. 7e, ‘fast’; see also grouped quantification in Fig. 7f). These results suggest that BLB treatment significantly increased the mobility of these long-range retrograde CTB-carriers by increasing the number of fast-moving carriers at the expense of the pausing and slow-moving ones.

We next further investigated the effect of radial contractility on the transport directionality of individual CTB-carriers within the microfluidic channels (Fig. 7a). We noted that their trajectories were predominately composed of two states: (i) a directed state and (ii) an undirected state, as indicated by the sloped and the vertical lines, respectively, in the displacement-time plot (Fig. 7h). Following 60 min blebbistatin treatment, the speed of the directed state was significantly increased, as indicated by the flatter slopes (Fig. 7h), whereas in the undirected state we observed pronounced back-and-forth motion (Fig. 7h, asterisks; sVideo 9), which has been previously described as low-efficiency trafficking pattern for long-range cargo transport (Yi et al., 2011). This is consistent with our observation that the ratio of the fast-moving CTB-carriers increases at the expense of slow-moving ones (Fig.7 e, f; see also sFig. 7k). To further quantify these back-and-forth movements, we compared the ratio of direction swap (*Sr*) in these tracks by measuring the ratio of the time cargoes spent travelling in the reverse direction (*t_rev_*) in relation to the total time travelled (*t_total_*), as shown in equate (1), with k being the number of trajectories.

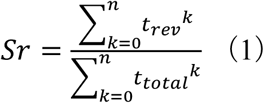

This analysis revealed that blebbistatin treatment significantly increased the amount of time cargoes underwent reverse motion, thereby increasing the ratio of direction swap in cargoes moving along the axon (Fig. 7i). Accordingly, we found that blebbistatin treatment decreased the number of CTB-positive carriers that traversed the imaging window within a given time (Fig. 7j), suggesting that the overall retrograde trafficking efficiency was reduced. These results support the positive role of axonal radial contractility in maintaining near-unidirectional retrograde trafficking, thereby ensuring the overall efficiency of long-range retrograde transport.

We next investigated how the radial contractility impacted the mobility of the directed and undirected carriers, respectively, by analyzing the dynamics of the CTB-positive carrier movements. To objectively and quantitatively analyze the effect of the contractility on the two motion states, we employed a two-state hidden Markov model (HMM) to annotate these CTB trajectories into undirected (D) and directed (DV) states (Fig. 7k), as previously described (Joensuu et al., 2017). The separating efficacy of this model was demonstrated by the fact that the step size of the undirected (0.1168 ± 0.45 µm) and directed transport states (0.4352 ± 1.53 µm) were distinct from each other. We then examined the effect of blebbistatin treatment, and found that it significantly increased the step size of large CTB-positive carriers in the DV state (Fig. 7l, pink spots), which is in good agreement with our earlier observation of the increased average (Fig. 7b) and instant speed (Fig. 7e-g; sVideo 9) of CTB carriers following short-term Blebbistatin treatment. For the D state in pretreated axons, CTB-positive carriers exhibited a much smaller step size (Fig. 7l), indicating that the mobility of these undirected carriers was constrained. However, this limited step size was significantly increased following 60 min blebbistatin treatment (Fig. 7l, blue spots). Taken together, the reduced pausing ratio (Fig. 7g) and an increased back-and-forth movement (Fig. 7h, i) suggests that the mobility of the undirected CTB carriers is increased following disruption of NM-II activity by blebbistatin. These results suggest the effect of 60 min blebbistatin treatment significantly increased the speed of both directed- and undirected-moving CTB carriers. Our findings indicate that the axonal actomyosin network maintains radial constriction, which not only impedes the speed of the directed fast-moving state but also suppresses the low-efficiency back-and-forth movement during the undirected state of these long-range carriers. The overall impact of this contractility on long-range trafficking is therefore positive, which facilitates the uni-directionality and the overall efficiency of long-range retrograde carriers.

### Prolonged inactivation of actomyosin-II disrupts the periodic actin rings and causes focal axon swelling (FAS)

We also used genetic approaches to explore the impact of longer-term manipulations of NM-II expression and activity on axonal structural stability. We used the commercially available short-hairpin RNA against MHC of NM-IIB (shNM-II) to specifically down-regulate the NM-II level in cultured cells. We found the endogenous NM-II level in transfected cells was significantly reduced (Fig. 8a-c). We also noticed a significant increase in the formation of FAS along the axons of shNM-II transfected neurons (Fig. 8b, arrowheads; Fig. 8d). We further used SiR-Actin to label the periodic actin rings in NM-II knock down neurons, and found that the periodic actin rings were largely disrupted. We detected accumulated actin blobs (Fig. 8e, arrowheads; Fig. 8f) and black-patches devoid of actin (Fig. 8e, brackets). As disassembly of actin MPS is one of the earliest steps underlying axon degeneration (Unsain et al., 2018; Wang et al., 2019), and FAS is a hallmark of irreversible axonal damage (Maia et al., 2015), our results suggested that disrupting axonal radial contractility could disturb the axonal MPS, leading to FAS and subsequent degeneration.

**Figure 8.**
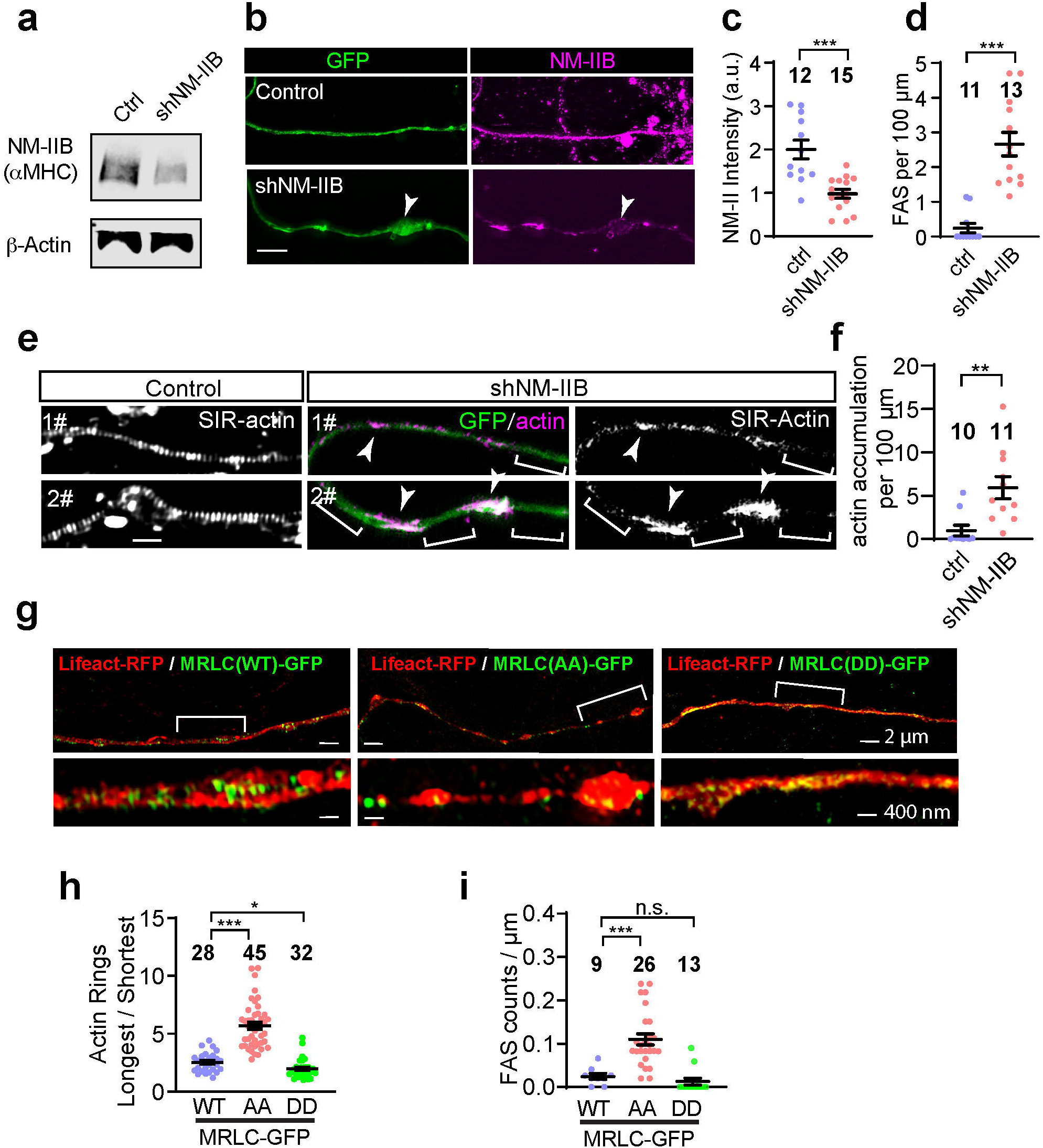
Long-term down-regulation of actomyosin-II activity disrupts the periodic actin rings and causes the formation of focal axonal swellings (FAS). **(a)** Knockdown efficiency of shNM-II construct. PC12 cells transfected for 48h with 2 μg of shRNA targeting MHC of NM-IIB (cat#:4390771; Invitrogen). Cell lysates were used to detect the endogenous NM-IIB levels by western blot. **(b)** Representative images of axons in shNM-II transfected neurons, arrowheads indicate the FAS. Bar=5 μm. **(c-d)** Quantification of the NM-II fluorescence intensity **(c)** and FAS number **(d)** of shNM-II axons shown in **(b)**. **(e)** Representative images showing that in the shNM-II axons, periodic actin structures were disrupted. The unevenly distributed F-actin-accumulation and F-actin-absent black patches were marked out with arrowheads and brackets, respectively. Bar=1 μm. **(f)** Quantification of the F-actin accumulation along the shNM-II axons. **(g)** Rat hippocampal neurons were co-transfected on DIV12 with the Lifeact-mRFP and either myosin-II regulatory light chain (MRLC) wild-type (MRLC (WT)-GFP), S19AT18A (MRLC (AA)-GFP), or S19DT18D (MRLC (DD)-GFP) plasmids. On DIV14 actin ring diameter fluctuations **(h)** and the FAS numbers per μm **(i)** of transfected axons were quantified. Data represent mean ± s.e.m; N values were shown in panels, representing the number of axons analysed; Data were from 2 independent preparations (two-tailed unpaired t-test, \**p*<0.05, \*\**p*<0.01, \*\*\**p*<0.001, n.s. no significant difference).

In addition to shRNA knock-down of endogenous NM-II, we also manipulated the NM-II activity by transfecting cultured neurons with the MRLC mutants, including S19AT18A (AA) or S19AT18A (DD) mutations, which abolish or enhance the NM-II activity (Beach et al., 2011), respectively. We found that 48 hours after NM-II (AA)-GFP transfection, the structural integrity of transfected axons was significantly disrupted as reflected by the significantly increased diameter fluctuations (Fig. 8g, h), abnormal accumulation of FAS (Fig. 8i) compared to those transfected with wild-type (MRLC(wt)-GFP) or constitutive active (MRLC(DD)-GFP) constructs. These results suggest that the long-term inactivation of actomyosin-II impairs the underlying periodic actin rings and causes irreversible structural damage to the axon, which eventually lead to degeneration.

## Discussion

Many factors affect long-range axonal cargo trafficking, including the number and type of attached molecular motors (Hancock, 2014; Vale et al., 1992), the polarity of the microtubule tracks (Kapitein and Hoogenraad, 2011), and friction from other organelles (Che et al., 2016). With the development of live-imaging microscopy, most of these factors have been extensively studied in cell-free *in vitro* systems (Vale et al., 1992), and within the axons of cultured neurons (Chowdary et al., 2015a; Chowdary et al., 2015b) and live animals (Bilsland et al., 2010). However, the impact of the narrow and rigid axonal plasma membrane on the transiting cargoes remains largely elusive. In this study, we have demonstrated that the transport of large membrane-bound cargoes causes an acute, albeit transient, radial stretching of the axonal plasma membrane, which is immediately restored by constitutive constricting forces generated by the membrane-associated periodic actomyosin-II network. We have also identified NM-II as a critical regulator of this radial contractility, which controls not only the speed but also the directionality of long-range cargo trafficking along the axon. Inactivation of this contractility machinery disturbs the axonal MPS, eventually generating FAS, which are early signs of degeneration. Our study therefore reveals novel functions of the actomyosin-II network underlying the structural plasticity of axonal MPS, which are not only required to facilitate the long-range axonal trafficking, but also maintain the structural stability of CNS axons.

### Radial contractility facilitates the overall efficiency of long-range retrograde axonal trafficking

An efficient long-range retrograde cargo transport machinery is critical for the survival and function of neurons (Barford et al., 2017; Bilsland et al., 2010; Tojima and Kamiguchi, 2015). Retrograde trafficking is driven by cytoplasmic dynein (Gennerich et al., 2007), which causes the near-unidirectional retrograde transport of nerve terminal-derived signalling endosomes and autophagosomes to the cell body (Chowdary et al., 2015b; Wang et al., 2015). Detailed analysis of the movement of these carriers has revealed that they are mainly composed of fast-moving retrograde-directed and undirected stalled carriers, with less than 3% being reverse-directed (anterograde) carriers (Wang et al., 2016; Wang et al., 2015). A similar near unidirectional motion pattern is shared by retrograde endosomes carrying nerve growth factor (NGF) (Chowdary et al., 2015b) and tetanus toxins (Lalli et al., 2003). The retrograde directionality of the fast-moving cargoes is driven by the progressive minus-end-directed dynein steps, the directionality of which is dependent on the opposing forces they receive (Gennerich et al., 2007). Actomyosin-II controls radial contractility, which poses a steric hindrance to the passaging cargoes (Che et al., 2016) and therefore could potentially affect the opposing force to their driven dyneins. However, the effect of the axonal actomyosin network on the dynein-driven trafficking is poorly understood. Early studies provided evidence that disrupting F-actin in axons does not interfere with organelle transport, which continues unabated or at an even faster rate (Morris and Hollenbeck, 1995), suggesting that the axonal F-actin network acts as a physical impediment to cargo transport. In line with this, we found that short-term blebbistatin treatment released subcellular tension, causing an expansion of the axon diameter and specifically increasing the transport speed of large cargoes. Our results therefore confirm that the directed fast-moving state of large cargoes is subjected to a constant impediment from radial axonal constriction. On the other hand, the undirected stalling of retrograde carriers is likely to be caused by either a balanced tug-of-war between kinesin and dynein (Belyy et al., 2016), or by the transient detachment of dynein-driven carriers from the microtubule tracks (Gennerich et al., 2007) which is known to be regulated by the load of the cargo (Yi et al., 2011). Our data using HMM-Bayesian partition show an increased mobility of undirected carriers after blebbistatin treatment, which suggests that axonal radial constriction might affect the microtubule attachment/tethering of these stalled carriers within the axon. This could be due to the tension-dependent tethering of dynein to the microtubule tracks (Cleary et al., 2014). Collectively, our data suggest that the force exerted by the plasma membrane on the carriers might also be important for the regulation of dynein-mediated transport. However, further study on the coordination between cargo trafficking and local axon radial tension, using higher temporal resolution live-imaging techniques, is needed to establish the precise relationship between axonal radial constriction and the motion of variously sized retrograde carriers.

### Radial contractility along the axonal shafts

The diameter of the long and thin axon has long been believed to be uniform for the same type of neurons. However, with the development of 3D EM reconstruction, diameter fluctuations have been detected along the length of axons in optical nerves (Giacci et al., 2018). Similarly, in live rat brains, axonal diameter fluctuations were revealed with super-resolution microscopy after the conduction of action potentials (Chereau et al., 2017). Consistent with these *in vivo* studies, we have used EM, SR-SIM and bright-field confocal microscopy to reveal that axons indeed undergo dynamic diameter fluctuations in cultured neurons. Using time-lapse SR-SIM to capture the dynamic changes of sub-diffractional membrane-associated structures in live axons, we directly visualized the dynamic radial expansion of both the axonal plasma membrane and of the periodic actin rings triggered by the passages of large axonal cargoes. These transient cargo-induced deformations indicate that: (1) the axonal plasma membrane exerts a constant tension to restrict its diameter; (2) the underneath periodic actin rings are constitutively constricted and can undergo radial expansion.

Indeed, these notions are supported with observations from previous studies. Zhang *et al*. have shown that the axon is the most rigid part of the neuron and is under constitutive tension (Zhang et al., 2017) from the ordered periodic longitudinal MPS composed of actin, spectrin, adducin and associated proteins (Xu et al., 2013). Depletion of adducin, which caps the barbed actin filaments in the periodic rings, causes progressive axonal dilation and degeneration (Leite et al., 2016), whereas the radial contractility remains unaltered, suggesting the existence of an alternative mechanism for axonal contractility. Moreover, axons are capable of sensing mechanical stimuli and rapidly change their diameter within seconds, by forming reversible axonal varicosities along their shafts (Gu et al., 2017). These plastic axonal deformations also point to the existence of regulative subcortical machinery, which might also involve the activity of actomyosin complexes. However, more live-cell studies using super-resolution microscopy are required to further examine how external factors can regulate axonal radial contractility in health and disease.

### Actomyosin-II complex is the structural basis underlying the axonal radial contractility

In mammalian cells, radial contractility of subcortical actin (Duan et al., 2018) or actin rings (Gormal et al., 2015) is regulated by mechanosensory NM-II, which has the ability to convert ATP into mechanical force. Recently, the activated form of NM-II light chain (p-MRLC) was shown to distribute with a similar periodicity and largely overlap with the actin MPS at the AIS (Berger et al., 2018). Moreover, depolarization rapidly decreased NM-II activity, indicating that this constricting structure is highly dynamic (Berger et al., 2018; Evans et al., 2017). Tropomyosin isoform Tpm 3.1, which activates and promotes the actin binding to NM-II, has recently been found to display a periodic distribution associated with subcortical actin rings in the AIS (Abouelezz et al., 2019b). In our study, we revealed that the ∼200 nm periodic actomyosin-II filaments are closely associated with actin MPS, in an NM-II-activity-dependent manner, suggesting the actomyosin structure indeed controls the contractility of periodic actin rings. However, considering that the length of NM-II filament is ∼300 nm (Hu et al., 2017), it is challenging to place it within individual periodic actin ring. To address this issue, we used triple-colour SIM labelling to visualize the localization of either MHC head domain or rod domain with actin rings. Our results point to NM-II head domain rather than the rod domains colocalizing more extensively with periodic actin rings, suggesting NM-II filament is likely to slide cross adjacent actin rings (intra-ring). This notion was further supported by our observation that manipulations of NM-II activity significantly altered the tilting angles of periodic actin rings. Indeed, the fact that NM-II is the force-generator that causes the constant contraction and deformation of the cortical actin filaments (Arnold and Gallo, 2014; Beach et al., 2011; Hu et al., 2017) is in good agreement with the recently identified role of actomyosin in controlling the dynamic structural plasticity in AIS (Abouelezz et al., 2019b; Berger et al., 2018; Evans et al., 2017) and long axon shafts (Chereau et al., 2017; Gu et al., 2017). Therefore, our results demonstrate that actomyosin periodicity goes far beyond the AIS, and serves as the structural basis for axonal radial contractility.

### Radial contractility maintains the axonal structural stability

Over-expansion of axonal segments, such as FAS, has been noted in several neurodegenerative diseases as well in traumatic brain injury, with the accumulation of organelles and cargoes at the axonal swellings - the presence of FAS is generally regarded as an early sign of axonal degeneration (Maia et al., 2015). We found that NM-II knock-down and MRLC inactive mutant expressing axons exhibited FAS, cargo accumulation and obvious signs of degeneration. These phenotypes were also observed in axons of adducin knockout mice (Leite et al., 2016). This strongly suggests that actomyosin-II-dependent radial contractility is critical to maintain the structural stability of the axon. Indeed, activity of NM-II motors is critical for F-actin turnover (Shutova et al., 2012), increasing the nucleation of actin filaments/bundles *in vitro* (Ideses et al., 2013), and promoting the fibre assembly of contractile ring and stress fibre in live cells (Abouelezz et al., 2019a; Tojkander et al., 2011). Disruption of the actin MPS is one of the earliest signs and likely the causal factor of axon degeneration, which is marked by FAS and fragmentation (Huang et al., 2017; Unsain et al., 2018; Wang et al., 2019). Given these facts, radial contractility is likely to be of great importance not only to ensure the stability of axonal MPS but also to maintain axonal structural integrity. Additional efforts combining live super-resolution imaging with computational and biophysical approaches will be needed to provide more insights into how axonal contractility is coordinated or disrupted under physiological and pathological changes in the CNS axons.

In summary, we have uncovered an inverse correlation between axonal cargo size and trafficking speed, and demonstrated that axons undergo transient deformation caused by the cargo transition in hippocampal axons. We have further identified the periodic structure of actomyosin-II along the axon shaft as the structural basis of axonal radial contractility. We have also characterized its role in facilitating long-range cargo trafficking by restricting inefficient back-and-forth cargo movement during the undirected state. Our data not only identify a novel role for the axonal actomyosin-II network in long-range cargo trafficking, but also highlight the importance of axonal membrane contractility in maintaining the stability of actin MPS along the axon.

## Material and methods

### Antibodies, molecular reagents and DNA constructs

SiR-Actin (#CY-SC001, Cytoskeleton); CellMask Deep Red (H32721; ThermoFisher Scientific); shRNA (#4390771; Invitrogen); p-MRLC (#3674; Cell Signalling); MRLC (#8505; Cell Signalling); Cell viability Kit (#L34973; ThermoFisher Scientific); Alexa-555- and Alexa-647-conjugated recombinant CTB were obtained from ThermoFisher Scientific (#c-34776, #c-34777). Mouse anti-synaptobrevin-2 (VAMP2) antibody was obtained from Synaptic Systems (#104211) and the rabbit anti-NM-IIB(ct) polyclonal antibodies were from Sigma-Aldrich (#M7939), mouse anti-NM-IIB(nt) monoclonal from Santa Cruz (sc-376954). Microspheres with fluorescence in all four channels were were purchased from ThermoFisher (TetraSpeck™; #7279). Alexa-647-phalloidin was purchased from Invitrogen (#A22287), while the mouse anti-β-tubulin III was from Covance (#MMS-435P). IRDye fluorescent secondary antibodies were from LI-COR (#925-32211; #926-32210); Alexa Fluor secondary antibodies were purchased from Life Technologies. The DNA construct encoding Lifeact-GFP was provided by Roland Wedlich Soldner (MPI Biochemistry, Martinsried), pTagRFP-mito was purchased from Evrogen (#FP147), pmRFP-LC3 was a gift from Tamotsu Yoshimori (Addgene plasmid # 21075). Lysotracker Red DND 99 (#7528; ThermoFisher). The remaining reagents were obtained from Electron Microscopy Sciences or Sigma Aldrich unless otherwise specified.

### Neuronal cultures

Hippocampal neurons were cultured from embryonic day 18 (E18) embryos from Sprague Dawley rats. All experiments were approved by The University of Queensland Animal Ethics Committee. Hippocampal neurons were prepared as described previously (Wang et al., 2015) and were plated on either glass coverslips (for confocal microscopy), plastic dishes (for EM) or in microfluidic chambers (Xona, #RD450) according to the manufacturer’s protocol (Taylor et al., 2005). For the pretreated groups, live-imaging of approximate 5 (30 12 μm) regions of interest (ROIs) was performed 2 h after the CTB labelling. For blebbistatin treatment, conditioned culture medium containing blebbistatin (10 μM) was only added to the middle and/or terminal chambers of the 6-well or 4-well microfluidic chambers (Xona, #TCND500; #RD450) to exclude its effect on the soma. For the short-term blebbistatin treatment, microfluidic devices were immediately returned to the 37°C imaging chamber for live imaging, and approximately 5 ROIs were imaged within a total duration of 60 min. For long-term blebbistatin treatment, microfluidic devices were returned to a 37°C CO_2_ incubator for an additional 2h before continuing the live imaging.

### Confocal microscopy

Stimulation and labelling were carried out on rat hippocampal neurons cultured in microfluidic chambers between day *in vitro* 14 (DIV14). Briefly, the culture medium was removed from all chambers and the neurons were incubated for 5 min at 37°C in labelling buffer (15 mM HEPES, 145 mM NaCl, 5.6 mM KCl, 2.2mM CaCl2, 0.5mM MgCl2, 5.6mM D-glucose, 0.5 mM ascorbic acid, 0.1% bovine serum albumin (BSA), pH 7.4), with 50 ng/ml CTB-Af555 or CTB-Af647 added to the nerve terminal chambers only. For Lysotracker labelling, the incubation time was 30 min. Neurons were then washed 3 times with warm neurobasal medium and returned to the original conditioned growth medium for 2 h prior to imaging. Images were acquired with a Zeiss LSM710 inverted microscope maintained at 37°C and 5% CO_2_, and movies were analysed for carrier kinetics using the spot function of Imaris software (Imaris7.7-9.2, Bitplane). Kymographs were generated using ImageJ software (NIH) using the plugin Multi-Kymograph for ImageJ. For immunofluorescence microscopy of fixed cells, the microfluidic devices were removed and neurons were subsequently fixed for 2-4 h at 4°C with phosphate buffered saline (PBS) containing 4% paraformaldehyde and 4% sucrose, followed by immunostaining as previously described (Wang et al., 2016). Permeabilization was performed using 0.1% saponin, 0.2% gelatin, and 1% BSA in PBS. Imaging was carried out on a Zeiss LSM710 confocal microscope and analysed with Zen (Zeiss) and ImageJ softwares. All images were compiled using Illustrator CS 5.1 (Adobe).

### Imaris tracing of axonal cargoes

Time-lapse movies of CTB-positive or Lysotracker-positive carriers were analysed for carrier kinetics using the spot function of Imaris software (Imaris7.7-9.2, Bitplane). In brief, region growth was enabled (threshold 50, diameter from border mode), estimated diameter 0.75 μm, tracing with autogressive motion (Max Distance 2 μm, Max Gap size 0). Resulted trajectories were filtered with duration > 10 s and instant speed > 0.07 μm/s. Average speed are calculated as track length divided by track duration. For Lysotracker-positive carriers that bleaches rapidly, only the diameter of the first time point in each trajectory were used as the diameter for size grouping.

### Co-labelling of F-actin and NM-II for SIM imaging

Cultured rat hippocampal neurons were fixed at DIV14. For dual-colour imaging using Phalloidin and NM-II, the fixation protocol was modified from that previously established for maintaining actin ultrastructure (Xu et al., 2013). Briefly, the samples were initially fixed in 4% paraformaldehyde dissolved in cytoskeleton buffer (CB, 10 mM MES, 150 mM NaCl, 5 mM EGTA, 5 mM glucose and 5 mM MgCl_2_, pH 6.1) for 30 min at room temperature and then blocked with antibody dilution buffer (2% BSA with 0.1% Triton X-100 in PBS) for 1 h at room temperature, after which the primary antibody (NM-IIB, diluted 1 in 500) and phalloidin-Af647 (0.14 µM) in 2% BSA in PBS were applied to the dish and incubated at 4°C overnight. Donkey anti-rabbit secondary antibody (Thermofisher, #A-21206) was diluted at 1/500 and incubated for 1h at room temperature. Samples were immediately mounted in Vectashield medium (Vector Laboratories, #H-1000) for SIM imaging. For the Triton X-100 extraction experiment, neurons were first treated with the extraction buffer (4% paraformaldehyde, 0.1% (v/v) Triton X-100, 1 μg/ml phalloidin in CB) for 45 s before the fixation and staining steps. For labelling of live neurons using SiR-Actin, newly dissolved SiR-Actin was added to culture medium at dilution rate of 1/1000, followed by incubation for 2 hour at 37°C. These neurons in glass-bottom dishes were then washed once with warm phenol-red free Neurobasal medium before imaging using ELYRA PS1 SR-SIM system (Zeiss) with at 30°C.

### Structured illumination microscopy

Imaging of live or fixed samples was performed using an ELYRA PS1 SR-SIM system equipped with a 100x objective (α Plan-Apochromat 100×/1.46 oil-immersion DIC M27) and a CMOS camera (PCO Scientific). For live-imaging of the SiR-actin labelled neurons, images were obtained with the Fastframe mode (100 ms exposure time, a time series of 200 frames at 20 s intervals, a SIM grating size of 51 μm at 647 nm wavelengths, and using 5 rotations). For fixed and stained samples, images were obtained by acquiring z-stacks of 10-16 slices with a spacing of 0.101 μm, an exposure time of 100 ms, a SIM grating size of 42-51 μm, and using five rotations. 3D structured illumination images were then aligned and processed using the Zen software. For cross-correlation analysis, line profiles were selected based on the standard of the existence of at least 4 consecutive NM-II peaks in a single axon shaft. The intensity profiles of each of the channels were then obtained using the Multichannel plot profile function of BAR collection (DOI10.5281/zenodo.28838) in ImageJ software (NIH). The auto-correlation or cross-correlation rate between the different channels was then examined using the xcorr function of Matlab. The correlation values for each axon segment were averaged and plotted.

### Western Blotting

Transfected PC12 cells were lysed with 2 x SDS loading buffer and homogenised with syringe and needle. The homogenate was boiled in 95°C for 5 min and fractionated by SDS-PAGE, then transferred to a polyvinylidene difluoride membrane. After blocked with Odyssey TBS blocking buffer for 30 min, membranes were washed once with TBST and incubate with antibodies against MRLC (1:1000), p-MRLC (1:1000), MHC (1:1000), or β-actin (1:5000) at 4°C overnight. Membranes were then washed three times with TBST and incubated with 1:20000 dilution of fluorescently tagged anti-mouse or anti-rabbit antibodies covered with foil for 1 h. Blots were washed three times with TBST and imaged with Odyssey imaging system according to the manufacturer’s protocols.

### Assessment of actin MPS and NM-II abundance

Periodic cytoskeletal structures (MPS) were defined as the axonal regions with at least 4 consecutive actin or NM-II peaks along the longitudinal direction. 5-7 of 5 µm × 5 µm ROIs were selected along the axons in each 3D-SIM image (50 µm × 50 µm). In each ROI, the length of axon with F-actin MPS was measured with ImageJ by a trained observer blind to the treatment conditions. In the same ROI, the particle number of NM-II staining was also automatically quantified with the Analyse Particle plugin of FIJI. MPS or NM-II abundance were then calculated as the percentage of segments length with an MPS or particle number over the total length of axons in the ROI respectively.

### Electron microscopy

Rat hippocampal neurons cultured in microfluidic devices (14-17 DIV) were treated as described for confocal microscopy (Wang et al., 2016), except that 10 µg/ml CTB-HRP was added to the nerve terminal chambers for the period of stimulation. Cells were returned to growth medium for 4 h prior to fixation. All cells were fixed in 2.5% glutaraldehyde for 24 h. Following fixation, they were processed for 3, 39-diaminobenzidine (DAB) cytochemistry using the standard protocol. Fixed cells were contrasted with 1% osmium tetroxide and 4% uranyl acetate prior to dehydration and embedding in LX-112 resin (Harper et al., 2011). Sections (∼50 nm) were cut using an ultramicrotome (UC64; Leica). To quantify CTB-HRP endocytosis, presynaptic regions were visualized at 60,000x using a transmission electron microscope (model 1011; JEOL) equipped with a Morada cooled CCD camera and the iTEM AnalySIS software. Membrane-bound compartments within the cell soma proximal region of the microfluidic channel were analysed, and the axon diameter measured using ImageJ software.

### HMM-Bayes analysis

Hidden Markov modelling (HMM) was used to predict the particle hidden states and the state transition probabilities from experimental trajectories. By using Bayesian model selection in the inference process, the simplest mobility model can be selected to describe these trajectories in an objective manner (Persson et al., 2013). We analysed the trajectories from each cell of interest using HMM-Bayes software (Monnier et al., 2015). A maximum of 2 hidden states was set to describe the trajectory movements, diffusion motion (D) and active transport state (DV), which were used to describe the undirected state and the directed state, respectively. In our cases, at least 10 channel ROIs were quantified for the control group and the blebbistatin-treated group, with corresponding trajectory numbers being 126 and 190 respectively. The D state with a low apparent diffusion coefficient state representing the immobile unattached movement. The DV state, which could be described by averaged velocity, represents the active transport attached movement. All of the analyses were performed using Matlab (R2016a, MathWorks, Inc.). The average step sizes of different transport states was calculated from all D-DV models.

### Statistics

We used GraphPad Prism 7 (GraphPad Inc.) for statistical analyses. Results are reported as mean ± s.e.m. For group comparisons, two-tailed nonparametric *t*-tests or paired *t*-tests were executed. *P* values < 0.05 indicated statistical significance. No statistical methods were used to predetermine sample sizes. Data distribution was assumed to be normal, but this was not formally tested. There was no formal randomization. Data collection and analysis were performed by different operators, who were blind to the conditions of the experiments.

## Supporting information

sVideo1

sVideo2

sVideo3

sVideo4

sVideo5

sVideo6

sVideo7

sVideo8

sVideo9

sVideo10

## Online Supplemental material

**s**Fig. 1 shows that the speed of Lysotracker-positive cargoes is inversely correlated with their size and is supporting Fig. 1. **sFig. 2** shows the axonal deformation caused by the unlabelled axonal cargoes in Lifeact-GFP expressing neurons and is supporting Fig. 3. **sFig. 3** shows the effects of NM-II short-term inactivation in actin MPS and is supporting Fig. 4. **sFig. 4** is an extended analysis on the contractility of actin MPS, supplementing Fig. 4. **sFig. 5** shows the colocalization between p-MRLC immunostaining and the actin MPS, supporting Fig. 5. **sFig. 6** shows the immunostaining of NM-II filaments and that of periodic actin rings, supplementing Fig. 1. 6. **sFig. 7** shows the inactivation of NM-II caused axon diameter expansion without affecting the microtubule structure or docking mitochondria, supplementing Fig. 7. The time-lapse images of axonal trafficking of CTB and Lysotracker in microfluidic devices are shown in **sVideo 1 and 2**. The time-lapse SIM showing the organelle-induced axonal expansions are shown in **sVideo 3**. **sVideo 4-5** showing the cargo induced plasma membrane deformation along the axon. **sVideo 6-7** showing the cargo-induced periodic actin ring expansion along the axon. **sVideo 8** showing the contractility of the periodic actin rings along the axon. The time-lapse images of axonal trafficking of CTB and Lysotracker in microfluidic devices perfused with Blebbistatin are shown in **sVideo 9 and 10.**

## Acknowledgements

This work was supported by grants from the Australian Research Council (ARC DE170100546 to TW) and the Australian National Health and Medical Research Council (GNT1120381 to FAM). FAM is an NHMRC Senior Research Fellow (GNT1060075). TW is an ARC DECRA Fellow (DE170100546). Imaging was performed at the Queensland Brain Institute’s Advanced Microscopy Facility, generously supported by the Australian Government through the ARC LIEF Grant (LE130100078 to FAM). Electron microscopy was performed at the Australian Microscopy and Microanalysis Facility, The University of Queensland. The authors would like to thank Rowan Tweedale and Tristan Wallis for editing the manuscript, and Joanne Jang, Nicholas Condon, Luke Hammond, Nick Valmas, He Huang, Xiaojun Yu, Victor Anggono and Rachel Gormal for their expert technical assistance. The authors declare no competing financial interests.

## Author contributions

FAM and TW designed the study, supervised the project and wrote the manuscript. TW performed live-imaging microscopy, confocal and SIM experiments and analysed data. WL performed HMM separation and assisted with data analysis. SM designed and performed EM experiments and helped with data analysis. AP designed and performed the analysis of particle speed and swap, and helped with SIM. JA performed the western blot experiment. AR and GS developed the staining and live-imaging protocol for dual-colour SIM. VL and PP helped with the edition of the manuscript and the figures. All authors discussed the results and commented on the manuscript.

## Abbreviations

BLB: Blebbistatin
CTB: Cholera toxin subunit B
DIV: Days *in vitro*
F-actin: Filamentous actin
FAS: Focal axonal swellings
NM-II: Non-muscle myosin II
ROI: Region of interest
MRLC: Myosin regulatory light chain
MHC: Myosin heavy chain

---

**Supplementary Figure 1.**
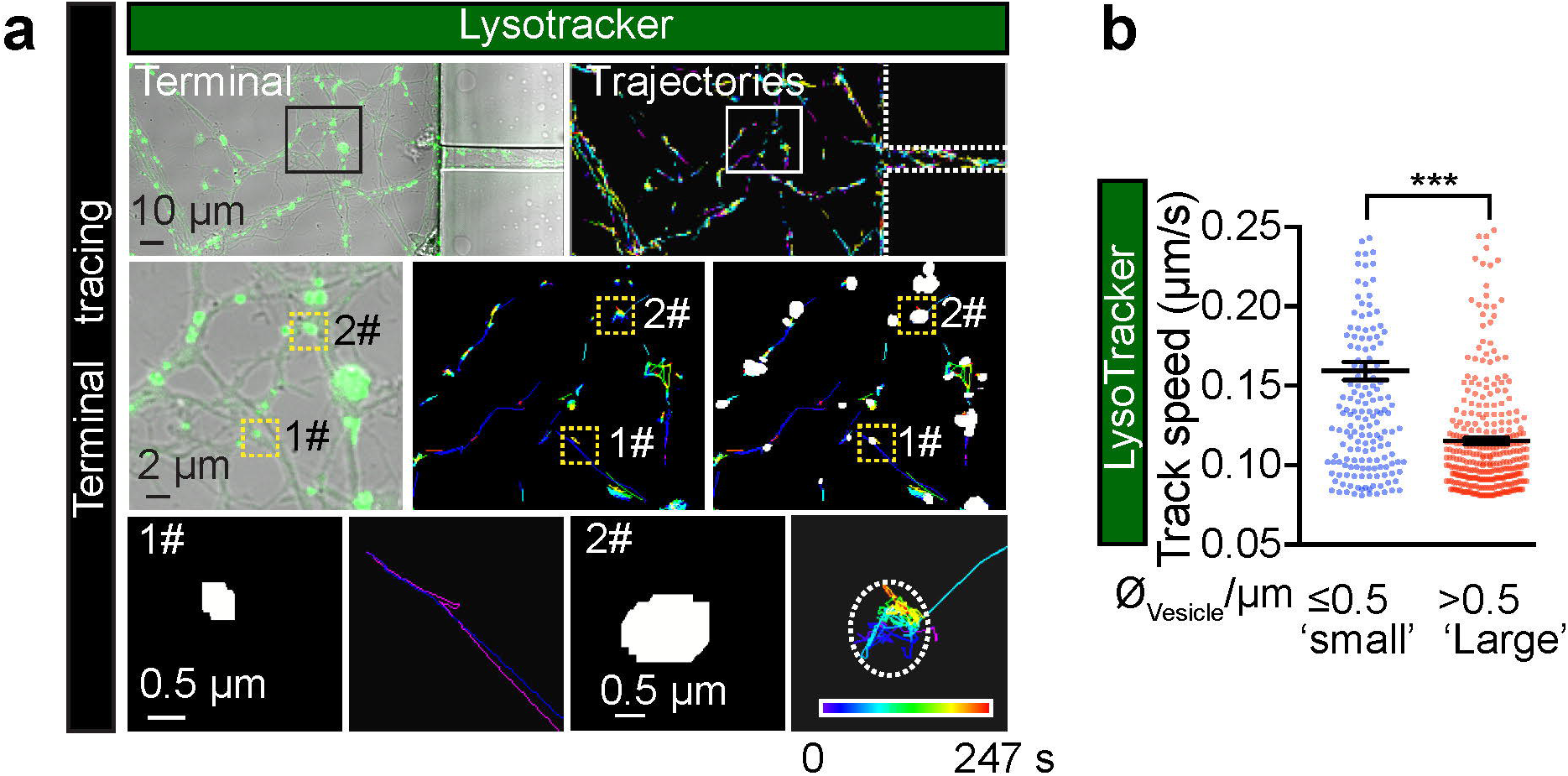
The speed of Lysotraker-positive cargoes is inversely correlated with their size in axons of cultured hippocampal neurons. **(a)** Representative time-lapse images of Lysotracker carriers at the nerve terminals of DIV 14 rat hippocampal neurons. Top-left: Lysotracker labelling at nerve terminals as isolated by the device. Top-right: Imaris tracing trajectories of the same region of interest. Trajectories of small (1#, diameter 0.5 μm) and large (2#, diameter > 0.5 μm) carriers were magnified in the bottom panels respectively. (**b)** Grouped analysis of average speeds of Lysotracker-positive cargoes with small (0.5 μm) and large (> 0.5 μm) diameters, showing a significant difference. Data represent mean ± s.e.m from 3 independent preparations (small, n=179, big n=342 tracks; two-tailed unpaired t-test, \*\*\**p*<0.001). The same data sets were also used in the pretreated group of Figure 7C.

**Supplementary Figure 2.**
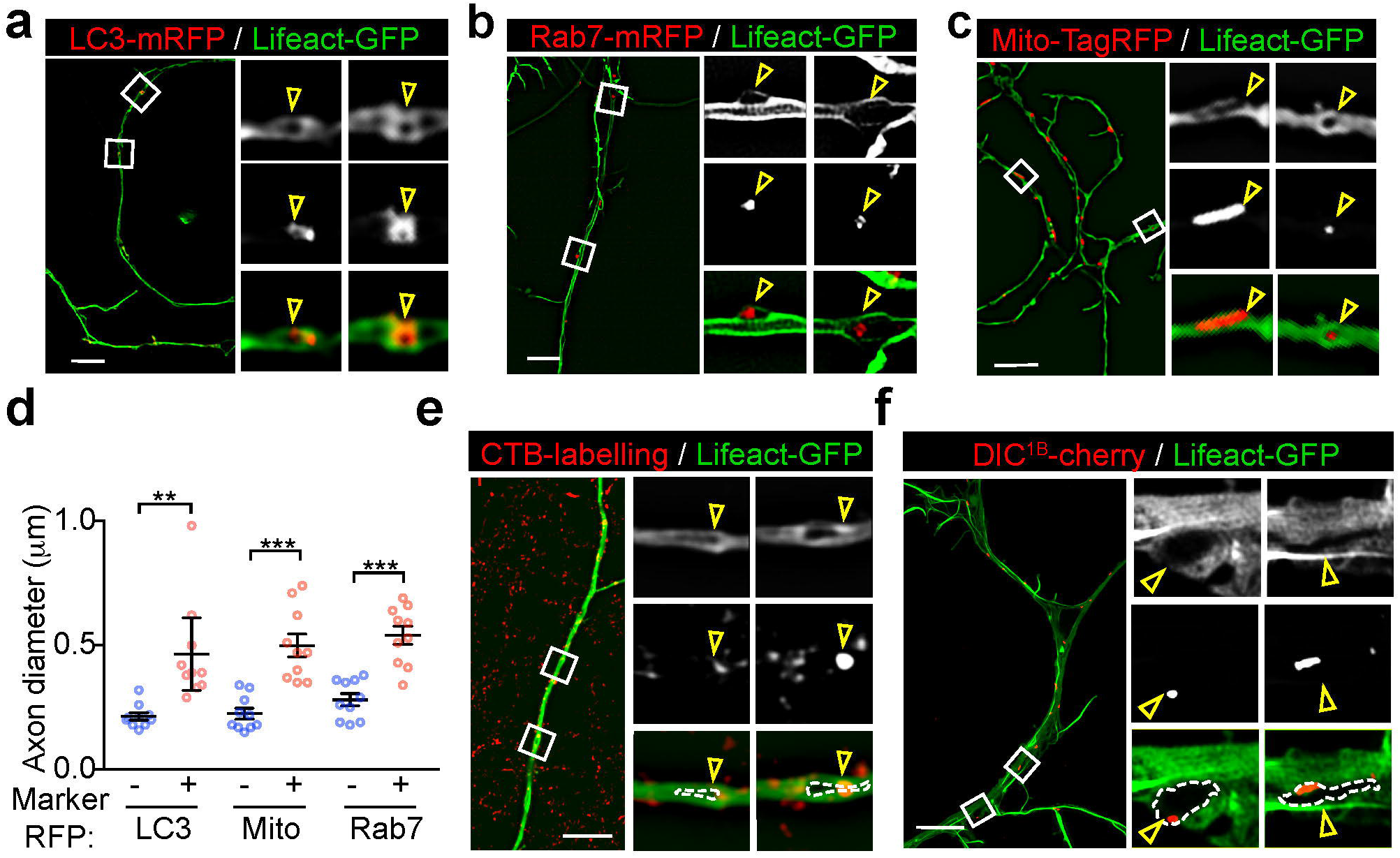
Large retrograde cargoes produce “fluorescence voids” within the axons of Lifeact-GFP-expressing neurons. Cultured hippocampal neurons grown in a microfluidic device were transfected on DIV12 with Lifeact-GFP and co-transfected with either LC3-mRFP (autophagosome) **(a)**, Rab7-mRFP (late endosome) **(b)** or Mito-TagRFP (mitochondria) **(c)**, and subjected to time-lapse imaging on DIV 14. Representative dual-colour 3D-SIM projections of neurons expressing Lifeact-GFP and with different subcellular markers are magnified in right panels, and overlapping regions are annotated. Bar = 5 μm. **(d)** Quantification of axon diameters with (+) and without (-) annotated markers. Data represent mean ± s.e.m (n=10 axons for each marker from 3 independent cultures; two-tailed unpaired t-test, \*\**p*<0.01; \*\*\**p*<0.001). **(e-f)** Representative dual-colour 3D-SIM projections of axons expressing Lifeact-GFP, which were co-labelled with the retrograde cargo marker CTB (**e)** or DIC^1B^ (**f)**. Boxed regions are magnified in right panels, overlapping regions are annotated. Bar = 2 μm **(e)** and 5 μm **(f)**.

**Supplementary Figure 3.**
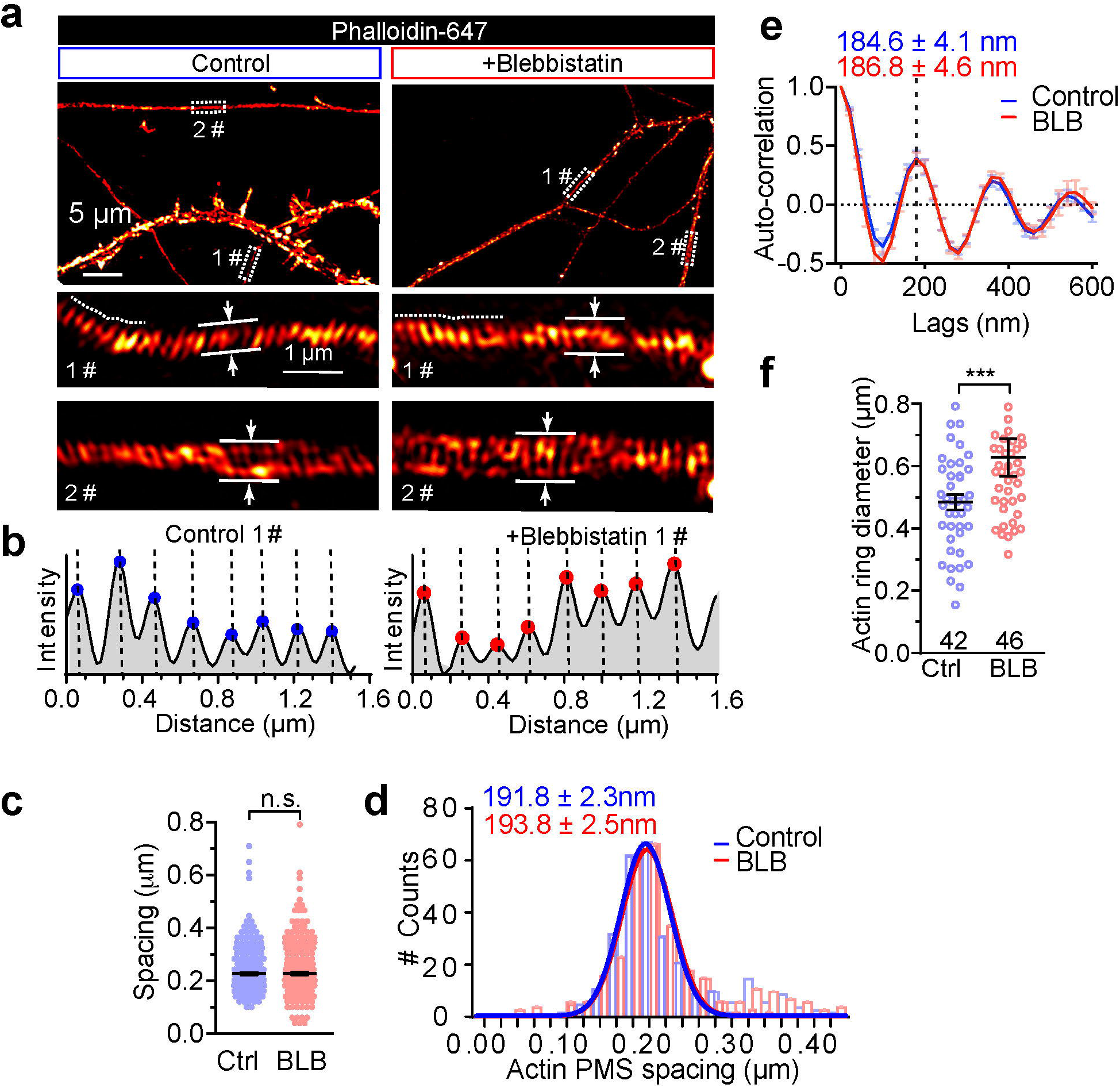
Short-term inactivation of NM-II increases the axon diameter without affecting the actin ring periodicity. **(a)** DIV14 rat axons were stained for endogenous F-actin (phalloidin) and imaged with 3D-SIM; two different boxed regions magnified with maximum z-projections shown in the lower panels. Axon diameters were measured as the average of a 1 µm segment. **(b)** Periodic actin peaks were identified using the ‘find peak’ function of BAR collection in Image J (SMA=1), along the line profiles as shown with dashed lines in 1# of **(a)**. x-value of the peaks were extracted, and the distance between adjacent peaks was shown in Fig. 5f, in both untreated (control) or short-term (60 min) blebbistatin-treated axons. **(c-d)** Comparison of spot plot **(c)** and Gaussian fitting curve **(d)** of periodic actin spacing distribution in control and blebbistatin-treated neurons. Data represent mean ± s.e.m, n=300 (Control), 316 (BLB) for periodicity quantification. **(e)** Autocorrelation analysis of the actin periodicity of control and blebbistatin-treated axons. Data represent mean ± s.e.m, n = 10 (Control) and 8 (BLB) axon segments were measured. **(f)** Quantification of actin diameters in control and blebbistatin-treated axons. Data represent mean ± s.e.m, n = 42 (Control) and 46 (BLB) actin rings diameters were measured. Values were measured from 3 independent cultures (two-tailed unpaired t-test, n.s. no significant difference; \*\**p*<0.05; \*\*\**p*<0.001).

**Supplementary Figure 4.**
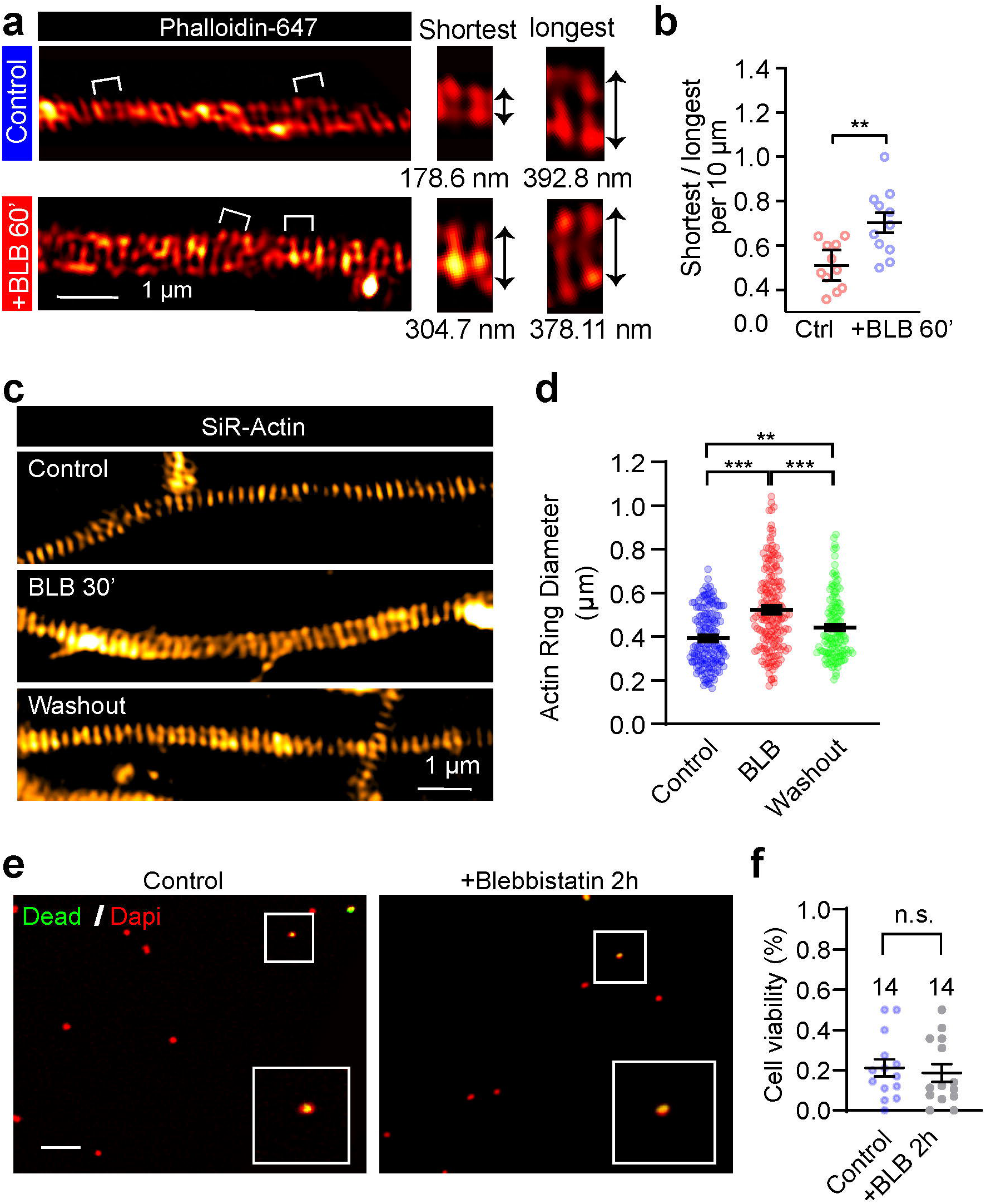
The effect of blebbistatin on periodic axon actin rings is reversible. (**d)** SIM images of endogenous F-actin (phalloidin) along the axon of a DIV14 rat hippocampal neuron before and after short-term blebbistatin treatment (10 µM, 60 min). Bracketed regions are magnified on the right, and the diameters of actin rings are shown below. (**b)** Quantification of actin ring diameter fluctuations; the diameters per 10 µm axon segments were measured and quantified. Data represent mean ± s.e.m, n=11 (Control) and 11 (BLB) axon segments were analysed. **(c)** In cultured hippocampal neurons, endogenous periodic axonal actin rings were labelled using SiR-Actin and live-imaged using SIM. Representative SIM images of axonal actin rings are shown of neurons following DMSO treatment (control), 30 min blebbistatin treatment (BLB), 12 h incubation after blebbistatin treatment and washout (Washout), respectively. Bar = 1 µm. **(d)** Quantification of diameters of the periodic actin rings along the axon. **(e)** Viability of neurons treated with 10 μM blebbistatin for 120 min. Boxed regions are amplified in the insets. Bar = 50 μm. **(f)** Quantification of viability rate. Data represent mean ± s.e.m. N values are labelled on the panels, representing numbers of axonal actin rings analysed. Values were measured from axons of at least independent cultures (two-tailed unpaired *t*-test, n.s. no significant difference; \*\**p*<0.01; \*\*\**p*<0.001).

**Supplementary Figure 5.**
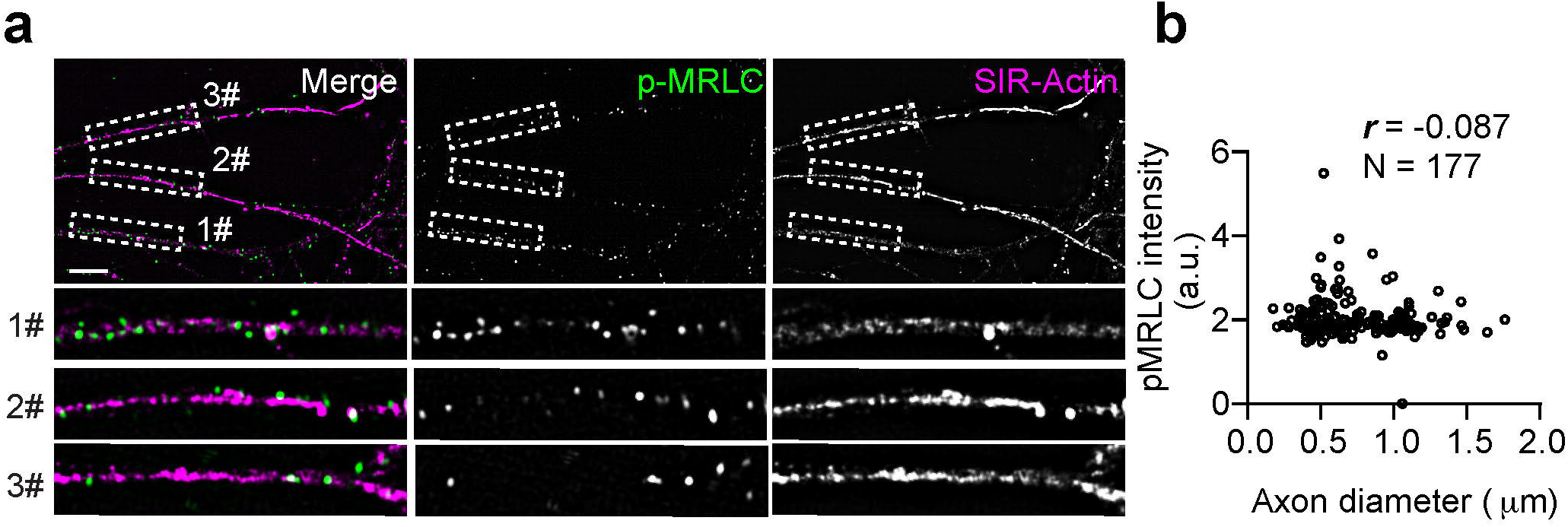
The phosphorylation of MRLC is not associated with the axonal diameter. **(a)** In DIV14 rat hippocampal neurons, endogenous periodic axonal actin rings were labelled using SiR-Actin followed by staining using diphosphorylated MRLC antibody (p-MRLC). Boxed regions are amplified in the bottom panels. Bar = 5 µm. **(b)** Pearson’s correlation coefficient (*r*) were calculated between the local p-MRLC level and the axon diameter. N values are labelled on the panels, representing numbers of axonal segments analysed. Values were measured from axons of 3 independent cultures.

**Supplementary Figure 6.**
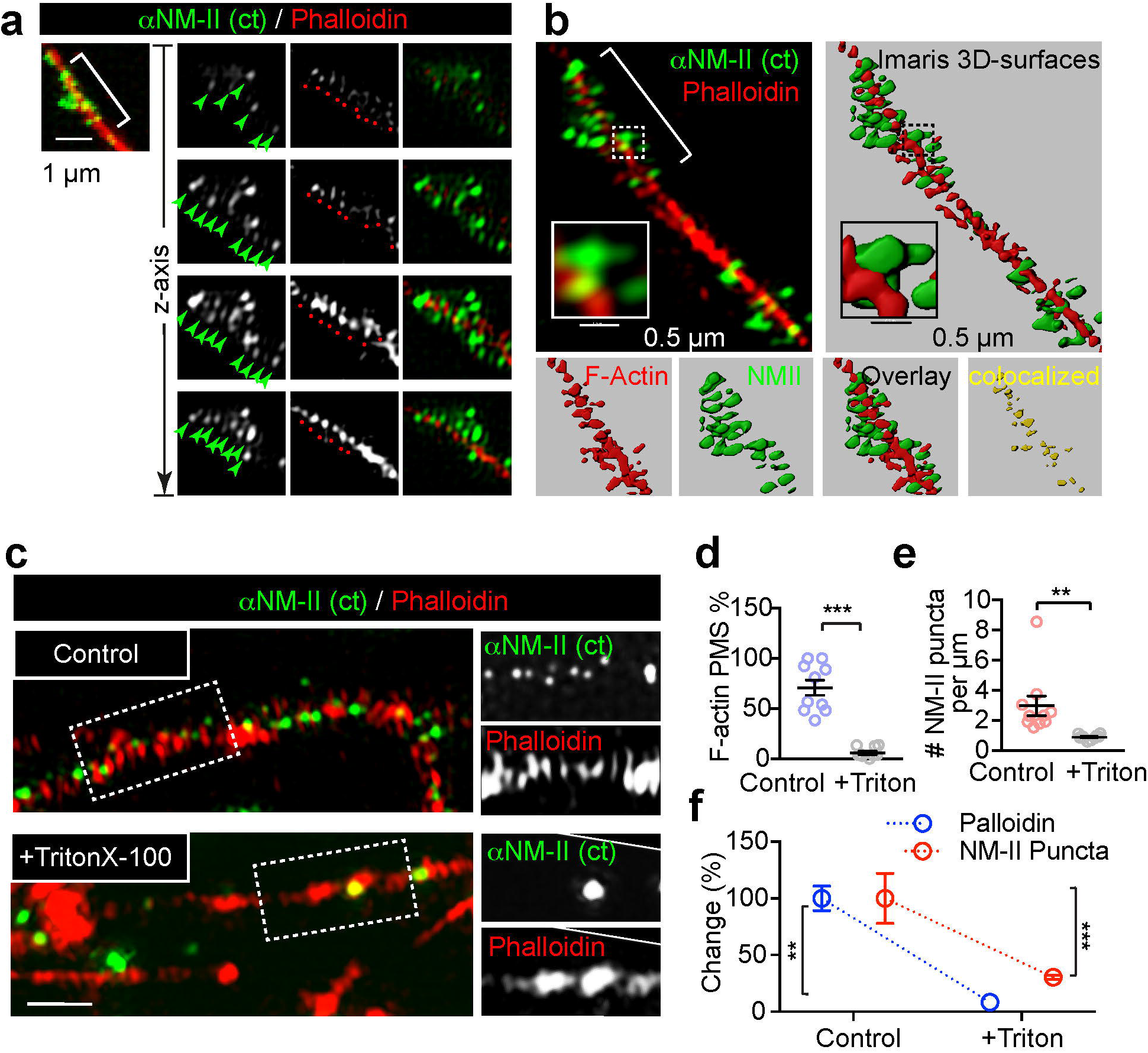
NM-II immunostaining closely correlates with the periodic actin rings along the axon. (**a)** DIV14 rat hippocampal neurons were stained for endogenous F-actin (Phalloidin) and NM-IIB and imaged with dual-colour 3D-SIM. Boxed regions are shown with individual z-stack planes, periodic structure of NM-II and actin are indicated with arrows (green) and spots (red) respectively. **(b)** The NM-II and actin structures resolved with 3D-SIM (right) were rendered into surface (right) using Imaris; boxed regions are magnified to show the accuracy of the rendering. The colocalization of the bracketed region is shown in the bottom panels. **(c)** Two-colour SIM images of endogenous F-actin (phalloidin) and NM-II along the axons of control and Triton X-100-extracted axons. Bar = 1 µm. (**d-f)** Quantification of the percentage of axons bearing periodic actin rings (**d)**, NM-II puncta number **(e)**, and the percentage change in these two parameters **(f)** in control and Triton X-100-extracted axons (+Triton). Data represent mean ± s.e.m, n=10 cells (Control) and 9 cells (Triton X-100) from 3 independent cultures (two-tailed unpaired t-test, \*\**p*<0.01; \*\*\**p*<0.001).

**Supplementary Figure 7.**
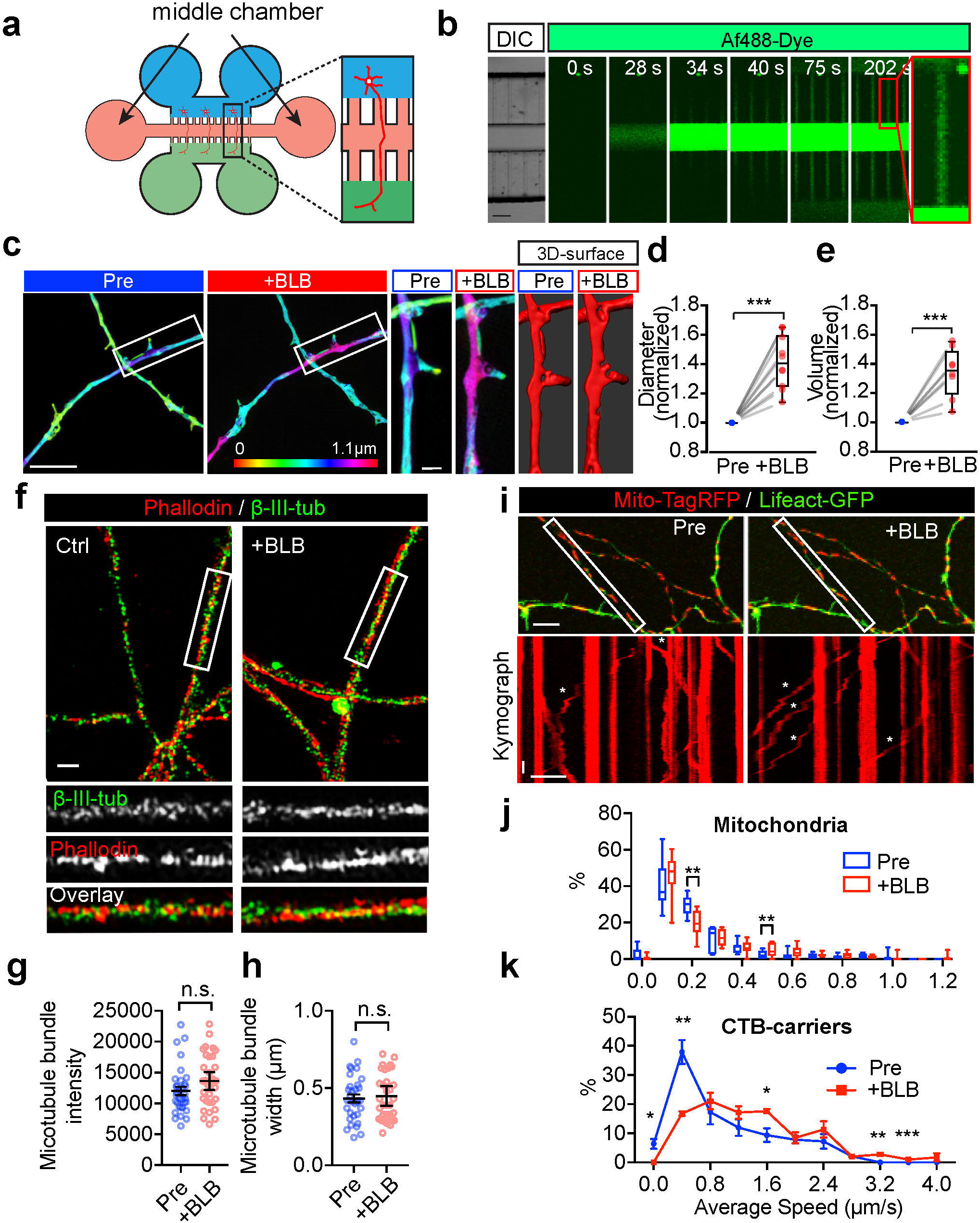
Short-term inactivation of NM-II causes the axon diameter expansion without affecting the microtubule structure or docking mitochondria. **(a)** Schematic cartoon showing the structure of a 6-well microfluidic device. **(b)** Time-lapse images showing the distribution of Af488 dye in the middle chamber of a 6-well microfluidic device, indicating its restriction capacity. Bar = 50 µm. **(c)** Axons of DIV 14 hippocampal neurons were treated with blebbistatin (10 μM) in the middle chamber for 90 min. Representative images of the 3D-SIM time-lapse images of axons before (Pre) and after blebbistatin treatment (+BLB). Value of z-axis is colour-coded. Scale Bar = 5 µm. Magnified images from the boxed regions and the corresponding Imaris surface rendered images are shown in panels on the right. Bar=1 µm. **(d-e)** Quantification of changes in the axon diameter **(d)** and volume **(e)** before and after blebbistatin treatment. Data represent mean ± s.e.m, for diameter quantification n=10 (Pre) and 10 (+BLB) axons were analysed, for volume quantification n = 8 axons (Pre), n = 8 (+BLB) axons were analysed, from 3 independent cultures (two-tailed paired t-test, \*\*\**p*<0.001). **(f)** DIV 14 hippocampal neurons were treated with blebbistatin (10 μM, 60 min), then fixed and stained for endogenous F-actin (phalloidin) and β-III-tubulin. Two-colour SIM was used to resolve the actomyosin and microtubule structures, respectively. Bar = 1 µm. Quantification of changes in the microtubule intensity **(g)** and bundle width **(h)** in control and blebbistatin-treated one. Data represent mean ± s.e.m, for intensity quantification n=33 (Pre) and 34 (+BLB), for bundle width quantification n = 33 axons (Pre), n = 37 axons (+BLB), from 3 independent cultures (two-tailed paired t-test, n.s. no significant difference). **(i)** Hippocampal neurons transfected with mito-TagRFP (red) and Lifeact-GFP (green) were imaged at the level of their axons before and after blebbistatin treatment. Kymograph of mitochondria movements from boxed regions are shown in lower panels. Asterisks indicate moving mitochondria. xBar=10 µm, yBar=10 s. **(j)** Quantification of average speed of mitochondria transport in **(i**). The frequency distribution of the trajectories with different speed are shown to note the effect on docking (0 and 0.1 μm/s) and moving mitochondria. Data represent mean ± s.e.m, n=7 (Pre) and 8 (+BLB) axons from 3 independent cultures (Student’s t-test, \**p*<0.05). **(k)** Quantification of average speed of CTB-positive carriers with and without blebbistatin treatment, as shown in transport in (Fig. 7a). The frequency distribution of the trajectories with different speed are shown to note the effect on slow and fast CTB-carriers, respectively. Data represent mean ± s.e.m, n=3 (Pre) and 3 (+BLB) experiments from 3 independent preparations (two-tail Student’s *t*-test, \**p*<0.05).

Supplementary Video 1. Retrograde trafficking of Lysotracker-labelled cargoes in the axon terminals.

The retrograde trafficking of axonal cargoes labelled with Lystotracker-deepRed in the terminal chamber of the microfluidic device. From top to bottom: The Lysotracker carriers overlapping with bright field signals and the masked carriers and trajectories are shown in the top panels, with the boxed regions of interest being magnified in the bottom panels. Bar = 5 µm.

Supplementary Video 2. Retrograde trafficking of CTB-labelled endosomes.

The retrograde trafficking flux of axonal cargoes labelled with CTB in an axon channel of a microfluidic device. From top to bottom: the CTB-labelled cargoes, their trajectories, and the trajectories that overlapped with the brightfield signals are shown, respectively. Bar = 5 µm.

Supplementary Video 3. The transit of cargo causes a transient radial expansion of the axon.

The passage of cargo-associated black-holes through the axon shafts caused an obvious mechanical stretching of the shafts, which are visualized in the Lifeact-GFP-expressing neuron (green) by time-lapse SIM. Black-holes that associated with moving cargoes are indicated with arrows. Bar = 2 µm.

Supplementary Video 4-5. Radial expansion of axonal actin rings caused by passages of lysosomal cargoes.

Time lapse dual colour SIM images at 20 s interval showing that diameter changes of periodic actin rings (SiR-Actin) correlate with the passage of large lysosomal cargoes (Lysotracker-Red). The bracket segment is amplified in the bottom panels. Diameter changes are indicated with arrows. Bar = 1 µm (top); 5 µm (bottom).

Supplementary Video 6-7. Radial expansion of axonal plasma membrane caused by passages of axonal cargoes.

Time lapse dual colour SIM images at 20 s interval showing that diameter changes of axonal plasma membrane (CellMask-Deep Red) correlate with the passage of large lysosomal cargoes (Lysotracker-Red). Diameter changes of axonal plasma membrane are indicated with arrows. Bar = 1 µm.

Supplementary Video 8. Dynamic radial contractility of periodic actin rings along axons.

Time-lapse 2D-SIM images at 20 s interval showing the diameter changes of periodic axonal actin rings periodic actin rings labelled with SiR-Actin. 4 bracket segments were amplified in the right panels. Diameter changes are indicated with arrows. Bar = 1 µm (left); 5 µm (right).

Supplementary Video 9. Retrograde trafficking of CTB-labelled endosomes with short-term blebbistatin treatment.

The retrograde trafficking flux of axonal cargoes labelled with CTB in an axon channel of a microfluidic device, after treatment with 10 µM blebbistatin for 60 min. From top to bottom: the CTB-labelled cargoes, their trajectories and the trajectories that overlapped with the brightfield signals were shown, respectively. Bar = 5 µm.

Supplementary Video 10. Retrograde trafficking of Lysotracker-labelled endosomes after blebbistatin treatment.

The retrograde trafficking of axonal cargoes labelled with Lystotracker in the terminal chamber of a microfluidic device, after treatment with 10 µM blebbistatin for 60 min. From top to bottom: the Lysotracker carriers overlapping with bright field signals and the masked carriers and trajectories are shown in the top panels, with the boxed regions of interest being magnified in the bottom panels. Bar = 5 µm.

